# Multiomic analysis of a pleural mesothelioma organoid biobank defines gene dependencies and therapeutic vulnerabilities

**DOI:** 10.64898/2026.11.16.766984

**Authors:** JA Valer, C Herranz-Ors, C Arnedo-Pac, M Shamseddin, J Obacz, SG Bhosle, M Garcia-Casado, S Brocklesby, J Cantwell, AE Andres, G Emanuelli, N Veale, J Espejo Valle-Inclan, CM Sauer, I Cortes-Ciriano, S Meakins, AE Beck, HE Francies, RC Rintoul, MJ Garnett, SJ Marciniak

## Abstract

Pleural mesothelioma (PM) remains an incurable cancer with few effective therapeutic options and limited understanding of its targetable dependencies, owing in part to a scarcity of clinically relevant, scalable models. Here, we describe establishment of an annotated biobank of 26 patient-derived PM organoids that faithfully recapitulate the clinical, genomic and histopathological landscapes of their parent tumours, including driver mutations, copy-number alterations and mutational signatures. By integrating these organoid models with a complementary collection of low-passage primary PM cell lines, we generate a functional precision oncology platform for PM. Genome-scale CRISPR-Cas9 screening across 23 patient-derived PM models uncovered 1,150 core fitness genes, including 638 not shared with pan-cancer core essentials, highlighting critical requirements for proteostasis, DNA repair, RNA processing and cell-cycle control. Parallel profiling of 106 compounds across 36 PM models revealed broad therapeutic vulnerabilities in proteostasis, growth signalling and epigenetic regulation. Beyond these shared vulnerabilities, both screens uncovered genotype-specific dependencies and drug responses linked to recurrent PM alterations. This functional map of shared and genotype-specific dependencies establishes patient-derived organoids as a scalable platform for precision therapeutic discovery in pleural mesothelioma.

## INTRODUCTION

Pleural mesothelioma (PM) is an asbestos-driven cancer and an archetypal occupational malignancy that remains among the most lethal of all solid tumours^1,2^. Contemporary series report a median overall survival of ∼12-18 months, with histology strongly shaping outcome (∼18 months for epithelioid versus ∼8 months for non-epithelioid disease)^3,4^. Treatment has changed little for decades, with platinum-pemetrexed chemotherapy anchoring systemic therapy, only recently being complemented by immune checkpoint blockade which has shown particular efficacy in non-epithelioid PM^3,5^. With a long latency between exposure and disease, PM will remain a substantial global health burden for decades to come. Incidence is projected to fall in countries where asbestos consumption peaked and was restricted earliest, but will keep rising in nations where use remains high and prohibitions on use have been delayed or are absent^6,7^.

The systematic molecular characterisation of large panels of cancer cell lines, combined with pharmacological and genetic perturbation screens, has enabled the construction of reference maps of cancer dependencies and therapeutic vulnerabilities that now are widely used for precision oncology^8–12^. However, PM is poorly represented in existing cell line collections, and available models capture only a narrow slice of the disease’s biological diversity^13,14^. Prolonged adaptation to conventional 2D culture can alter cellular phenotypes and gene-expression programmes, making it difficult to distinguish culture-induced effects from tumour-intrinsic biology. Furthermore, most established cell lines lack matched germline samples and comprehensive clinical annotation, limiting confident somatic variant interpretation and genotype-phenotype analyses^13^. Critically, long-term 2D culture can favour the expansion of selected cellular populations, reducing the heterogeneity present within the original tumour. In contrast, pleural mesothelioma often exhibits marked intra-tumour heterogeneity and subclonal diversity *in vivo*, features that are thought to influence therapeutic response and the emergence of treatment resistance^15,16^. These constraints impede mechanistic discovery and slow the identification of actionable vulnerabilities in a cancer with pressing unmet clinical need.

Tumour organoids, 3D cultures derived from patient tissue and propagated in extracellular matrix with defined niche factors, offer a route to models that better preserve tumour architecture, genotype and functional state than conventional 2D systems^13,14^. PM organoid protocols have used diverse derivation strategies, with variable success in long-term expansion and biobanking^17–19^. In contrast to the large, deeply characterised organoid biobanks available for some carcinomas^11,20,21^, PM organoid efforts have remained small, models are variably annotated, and poorly developed for the kinds of downstream functional genomics and screening workflows that demand robust growth, genetic tractability and experimental reproducibility at scale.

Here, we establish a renewable, deeply characterised biobank of PM organoids generated using an optimised, defined culture medium that enables derivation and long-term propagation directly from patient tumour material without pleural fluid supplementation. We pair biobanking with molecular annotation and show that this platform supports scalable functional interrogation, including whole-genome CRISPR screening and high-throughput drug profiling, enabling systematic mapping of mesothelioma dependencies and therapeutic vulnerabilities in a disease-relevant, patient-proximal model system.

## RESULTS

### A UK-wide pleural mesothelioma organoid biobank integrating multimodal profiling

We established a UK-wide acquisition and processing network for pleural mesothelioma biosamples coordinated through Mesobank UK, linking four contributing clinical sites with a centralised derivation and profiling pipeline (Fig. 1A). For each donor, fresh tumour tissue and, where available, pleural fluid, alongside matched peripheral blood for germline reference, were collected. Samples were transferred to a central facility for organoid derivation, where systematic protocol optimisation was performed. Culture conditions were benchmarked across 31 samples from 20 donors, comparing commercially available tumour and airway epithelial growth media with an in-house formulation adapted from published epithelial, lung, and airway organoid media, tested alone and with supplementation^22–26^. Derivation efficiency was highest with the in-house formulation, which was therefore selected for further refinement. The contribution of individual components was then mapped by systematic withdrawal and add-back across nine independent organoid models. This yielded a defined culture medium that enables derivation and long-term propagation directly from patient tumour material, without the need for pleural fluid supplementation (see Methods). Separate protocols were established for derivation from primary PM tissue and from pleural fluid, and refined until each met stringent criteria for derivation success (Supplementary Fig. S1A&B, Methods). A model was considered successfully derived if it expanded to more than 25 million cells, yielded a 25-cryovial bank, survived cryopreservation and freeze-thaw cycles, remained free of contamination and fibroblast overgrowth, and passed model-tumour concordance testing by whole-genome sequencing (WGS), including matched blood where available, and by immunohistochemistry (IHC), where parent tumour material was available. Successfully derived models underwent molecular, functional and histopathological characterisation and were linked to longitudinal clinical data, generating a clinically annotated resource of PM models (Fig. 1A).

**Fig. 1.**
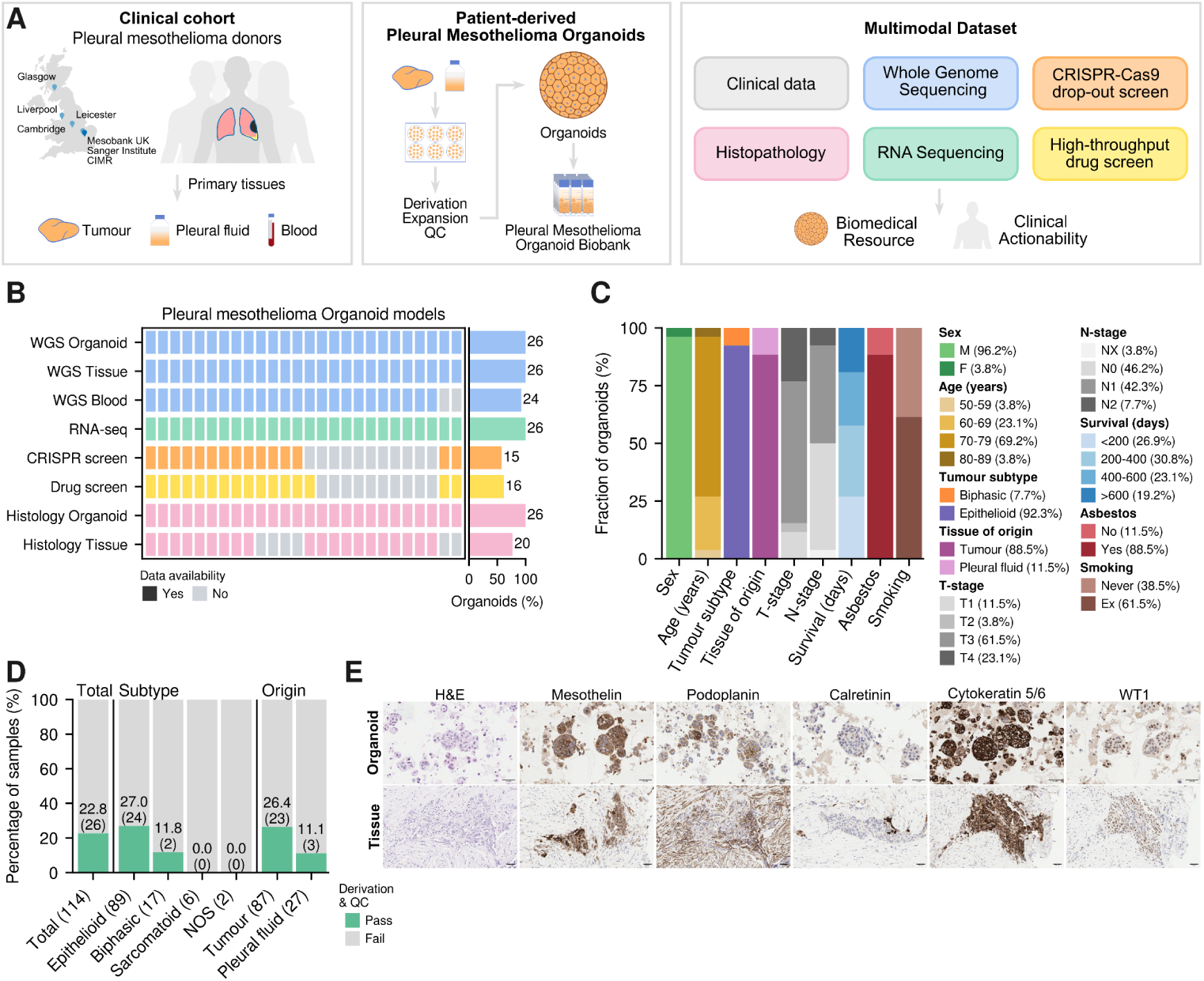
A pleural mesothelioma organoid biobank with linked multimodal profiling. (A) Overview of the contributing centres (Cambridge, Leicester, Liverpool and Glasgow) and the biospecimens collected (tumour tissue, pleural fluid when available, and matched blood), alongside schematics of the derivation workflow and the integrated dataset generated. (B) Data completeness across the biobank (organoids n = 26) for molecular profiling and functional screens. (C) Biobank cohort characteristics and sample attributes, including sex, age, histological subtype, tissue source for derivation, TN stage, survival from sampling, asbestos exposure and smoking status. (D) Derivation success rates stratified by histology and specimen source. Total attempted derivations are detailed in each label as (n), success rate and successfully banked organoids are detailed in each category as %(n). (E) Representative matched primary tumour and organoid sections (model WTSI-MESO_089) stained with H&E and immunohistochemistry for mesothelioma markers (mesothelin, podoplanin, calretinin, CK5/6 and WT1). Scale bars are 50 μm.

We generated a biobank of 26 patient-derived pleural mesothelioma organoid models with broad multimodal data completeness, representing the largest and most deeply characterised PM organoid resource to date (Fig. 1A&B). WGS was obtained for all organoids and their matched tumours, with matched germline available for 24 donors. RNA sequencing was performed for all models, while genome-scale CRISPR-Cas9 screens and high-throughput drug profiling were completed for substantial subsets (15 and 16 models, respectively). All models underwent histopathological assessment, and material from the parent tumour was available in 20 cases, allowing systematic comparison of tissue architecture and immunohistochemical marker expression between parent tumour and model (Fig. 1B, Supplementary Table 1).

The resulting cohort captured key clinicopathological features of pleural mesothelioma (Fig. 1C), including a predominance of older donors (median=74 years, IQR=71-76, range=54-80) and a male-biased sex distribution, consistent with the typical clinical presentation of PM^27^. Most tumours were classified as epithelioid, with a smaller proportion representing biphasic histology. Asbestos exposure was reported in most cases, alongside a mix of never- and ex-smokers. Survival intervals from time of sampling highlighted the aggressive natural history of this disease (median=300 days, IQR=191-545, range=64-992) (Fig. 1C, Supplementary Table 1). Organoid derivation efficiency varied by histological subtype and specimen source. Success rates were highest for epithelioid tumours, with primary tumour tissue yielding higher derivation efficiency than pleural fluid. These observations informed prioritisation of tumour tissue for subsequent derivations, while still supporting successful organoid generation from pleural fluid in selected cases (Fig. 1C&D, Supplementary Table 2).

In parallel, we performed a similar multimodal analysis on a set of previously established, low-passage primary PM 2D cultures^28^. Where available, the clinical features of the 2D models were comparable to those of the organoid cohort. WGS and RNA sequencing were obtained for all primary PM cell lines, while genome-scale CRISPR-Cas9 screens and high-throughput drug profiling were completed for substantial subsets (9 and 20 models, respectively) (Extended Fig. E1A-C, Supplementary Table 3).

To assess fidelity at the level of tissue architecture and lineage identity, we performed matched histopathological and immunohistochemical analyses between primary tumours and derived organoids (Fig. 1E and Supplementary Fig. S2). PM organoids formed compact, multicellular 3D structures embedded in basement membrane extract, recapitulating key aspects of tissue organisation. Across the cohort, organoids and parental tumours showed broadly concordant expression of established mesothelial markers, including mesothelin, podoplanin, calretinin, cytokeratin 5/6, and WT1, which are routinely used to distinguish mesothelioma from other thoracic malignancies^29^. As expected, marker expression varied across samples, reflecting biological heterogeneity within PM, and no single marker was universally expressed. Many markers are associated with epithelioid differentiation, and their absence in subsets of cells is consistent with known intra-tumoral diversity. Despite this, concordance between tumour and organoid was high, with preservation of both positive and negative staining patterns in most models. Observed differences between model and matched index samples were typically modest and can be explained by the pronounced spatial heterogeneity of primary tumours. Whole-slide sections frequently contained intermixed populations with distinct marker expression states (e.g., WTSI-MESO_086 and WTSI-MESO_008), and organoids may derive from one or multiple such subpopulations. Notably, a subset of organoids preserved this immunohistochemical heterogeneity, maintaining mixed marker-positive and marker-negative populations (e.g., WTSI-MESO_001) (Supplementary Fig. S2).

Using our newly-defined media conditions, these analyses demonstrate that patient-derived mesothelioma organoids preserve key histopathological and immunophenotypic features of their parental tumours while capturing aspects of intra-tumour heterogeneity. Together with their robust genomic and functional tractability, this supports the utility of our biobank as expandable, physiologically relevant models for systematic investigation of mesothelioma biology.

### Pleural mesothelioma organoids recapitulate the genomic landscape of matched tumours

We applied WGS to comprehensively characterise the genomic features of all PM models, assess the fidelity of our PM organoid biobank to matched primary tumours, and compare the cohort genomic profiles against the established genomic landscape of pleural mesothelioma^30–34^ (Fig. 2; Extended Fig. E2). Both parental PM tumours and derived organoids exhibited the characteristically low somatic mutation rate of PM, typically ≤2 single-nucleotide variants (SNV) and indel mutations per Mb^31^, and mutational burden was closely matched within each tumour–organoid pair (Fig. 2A; Supplementary Table 4). The fraction of shared mutations between matched tumour and organoid was high (median=72.4%, IQR=(64.0-79.6)%) (Extended Fig. E2A&B). Interestingly, our cohort captured one tumour-organoid pair (WTSI-MESO-098) with an elevated burden of both SNVs and indels, consistent with a tumour-intrinsic hypermutated state faithfully recapitulated in the organoid (Fig. 2A). This uncommon PM hypermutant case is further discussed later. Two additional models with increased mutation burden (WTSI-MESO-006 and 009), which were derived from tumour tissue and pleural fluid from the same donor, are explained by the lack of matched germline WGS. Ploidy estimates were similarly concordant between tumours and derived organoids, with most samples remaining near diploid (Fig. 2A; Supplementary Table 4), supporting preservation of large-scale genome architecture through derivation and expansion.

**Fig. 2.**
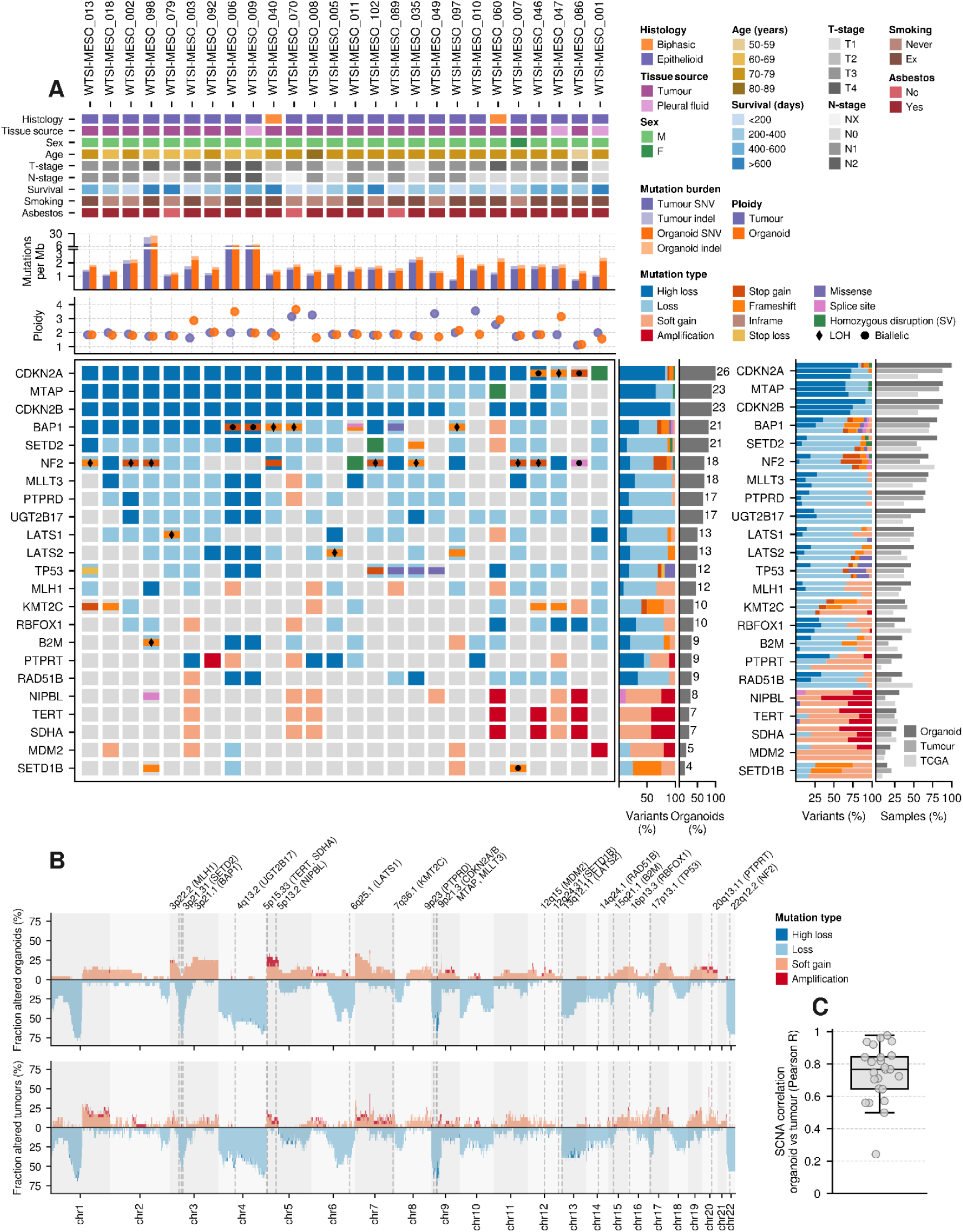
Whole-genome sequencing shows pleural mesothelioma organoids recapitulate the genomic landscape of matched tumours. (A) Per-model clinical annotation and tumour–organoid concordance for mutational burden and ploidy, together with the landscape of recurrent driver alterations. Driver alterations are shown with event type, alongside summary bar plots comparing alteration modes across organoids, matched tumours and TCGA mesothelioma samples (right). (B) Genome-wide somatic copy-number alteration (SCNA) profiles for organoids (top) and matched tumours (bottom), with key loci indicated (e.g., *MLH1*, *SETD2, BAP1* and others), illustrating preservation of hallmark pleural mesothelioma copy-number patterns. (C) Matched tumour–organoid SCNA concordance across the cohort (Pearson r), showing high genome-wide similarity (n=24). Each point is a tumour-organoid pair; the box indicates the median and interquartile range (IQR), whiskers extend to 1.5 × IQR.

Consistent with the known biology of PM, the driver landscape across organoids was dominated by tumour-suppressor inactivation and copy-number loss, and closely mirrored both matched tumours and TCGA PM samples^31^ (Fig. 2A&B; Extended Fig. E2A; Supplementary Table 5). *CDKN2A* was recurrently inactivated across donors, predominantly through copy-number loss, with frequent co-alteration of the adjacent *MTAP* and *CDKN2B* loci (Fig. 2A&B). Additional established PM drivers showed the expected mutational frequencies and alteration types, including *BAP1, NF2, SETD2, TP53, LATS1, LATS2* and *SETD1B* (Fig. 2A). For each gene, the relative contribution of particular mutational events (e.g., copy number loss) was highly concordant across organoids, matched tumours and the TCGA PM cohort (Fig. 2A; Extended Fig. E2A). Less frequent alterations included cancer driver genes previously reported in PM (e.g., RBFOX1, B2M and TERT) as well as additional candidates for follow-up in larger cohorts (Fig. 2A; Supplementary Table 5). Altogether, these results confirm that organoid derivation faithfully preserves the mutational spectrum of PM driver genes.

To evaluate copy-number architecture at genome wide resolution, we compared somatic copy-number alteration (SCNA) profiles for each organoid and its matched tumour (Fig. 2B&C; Supplementary Fig. S3). Organoid SCNAs recapitulated hallmark PM patterns, including recurrent loss of 1p, loss of 3p (encompassing regions containing *BAP1*, *SETD2* and *MLH1*), loss of chromosome 4, loss of 6q (including *LATS1*), focal/arm-level loss on 9p (including *CDKN2A/B* and *MTAP*), and loss of 22q (including *NF2*), together with recurrent gains such as 5p (including *TERT*), consistent with previously reported copy-number landscapes of pleural mesothelioma (Fig. 2B)^31,33^. Matched SCNA concordance between each organoid and its tumour was high (Pearson R median=0.77, IQR=0.65-0.84), indicating strong preservation of copy-number structure across organoid derivation and culture (Fig. 2C; Supplementary Fig. S3). In line with this, assessment of banked organoids together with their matched early-passage version confirmed the stability of the mutational landscape of point mutations and SCNAs (Supplementary Fig. S4A-C).

These data demonstrate that patient-derived PM organoids faithfully recapitulate the genomic mutational landscape and clonal composition of their parental tumours, while preserving tumour heterogeneity and enabling systematic, model-resolved comparisons across clinically annotated cases.

To compare these findings against alternative *in vitro* systems, we performed analogous genomic analyses on 21 low-passage PM cell lines (Supplementary Fig. S5). While broadly recapitulating the same spectrum of driver gene alterations and CNA patterns observed in organoids and primary tumours, primary PM cell lines showed higher mutational burden and greater variability in ploidy (Supplementary Fig. S5A&B; Supplementary Fig. S6; Supplementary Tables 4&6). Nevertheless, increased mutational burden in low-passage PM primary cell lines were modest compared to established PM cell lines (see Methods): PM organoids median=1.86, IQR=1.62-2.36; PM primary cell lines median=8.74, IQR=8.53-9.04; and established PM cell lines median=22.92, IQR=20.87-26.74 (Supplementary Fig. S5C). Clonal fractions were higher in primary PM cell lines (median=0.87, IQR=0.86-0.88) compared to those in primary tumours and derived organoids (tumours median=0.60, IQR=0.55-0.67; organoids median=0.65, IQR=0.58, 0.71), consistent with reduced heterogeneity in primary cell lines relative to organoids (Extended Fig. E2C; Supplementary Fig. S5D; Supplementary Table 4). These results suggest that organoids more closely preserve the genomic features of primary tumours.

### Mutational processes are preserved between pleural mesothelioma organoids and primary tumours

To assess whether patient-derived PM organoids faithfully recapitulate the mutational processes characteristic of primary tumours, we compared mutational burden, mutational profiles, and mutational signatures between organoids and matched tissues. Across samples with matched germline data, mutational burden was uniformly low (≤2 mutations/Mb) in both organoids and primary tumours, consistent with previous genomic studies demonstrating that PM is a genomically quiet tumour with a low mutational burden (Fig. 3A; Supplementary Table 4). This was accompanied by a predominance of SNVs over indels, in line with previous reports^33,34^.

**Figure 3.**
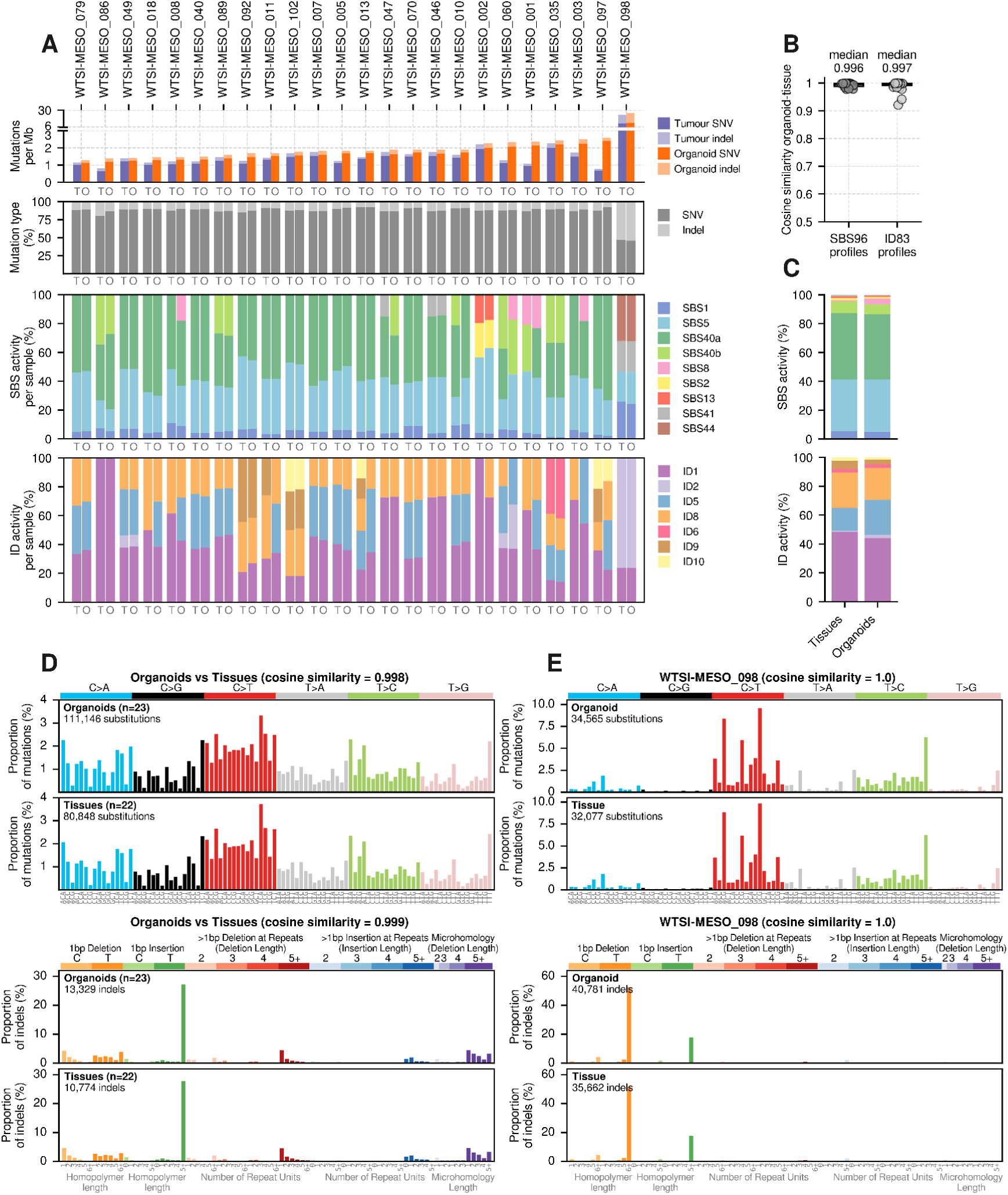
Preservation of mutational burden and mutational signatures between PM organoids and matched primary tumours. (A) Per-sample mutational burden and relative mutation type comparing matched tumour (T) and organoid (O) pairs, stratified by mutation class (SNVs vs indels). Bar plots additionally show relative contributions of single-base substitution (SBS) and indel (ID) mutational signature activities. (B) Cosine similarities between matched tumour–organoid pairs based on SBS96 and ID83 mutational profiles, demonstrating high concordance across all sample pairs (n = 24 pairs). (C) Aggregated relative contributions of SBS and ID mutational signature activities depicted in 3A. (D) Aggregate SBS and ID mutational profiles. Upper left: combined SBS profiles for non-hypermutant organoids (n=23) and matched tumours (n=22), showing high similarity. Lower left: corresponding indel profiles for organoids and matched tumours, showing high similarity. (E) Mutational profiles of the hypermutant case (WTSI-MESO_098). Upper right: SBS profiles for organoid and matched tumour. Lower right: indel profiles for organoid and matched tumour, demonstrating concordant hypermutator phenotype.

We next quantified the concordance of mutational profiles between organoids and matched tumours. Cosine similarity analysis demonstrated near-identical profiles for both SNVs and indels within each pair (median cosine similarity: 0.996 for SNVs; 0.997 for indels) (Fig. 3B; Supplementary Fig. S7&S8). This high degree of similarity was also maintained at the cohort level, where aggregated mutational profiles of organoids were nearly indistinguishable from matched tumours (cosine similarity: 0.998 for SNVs; 0.999 for indels) (Fig. 3D). These findings indicate that our organoid models robustly preserve the global mutational landscape of their parental tumours. Similarly, banked organoids and their matched early-passage versions presented near-identical profiles and signature activities, indicating that neither the mutational landscape nor the processes generating it are appreciably altered by culture. (Supplementary Fig. S4D-F).

To examine the underlying mutational processes, we performed *de novo* extraction of SNV and indel signatures followed by assignment to COSMIC reference signatures (Supplementary Table 7&8). Most organoid-tissue pairs exhibited activity of SBS1, SBS5, and SBS40, consistent with ubiquitous clock-like mutational processes associated with endogenous DNA damage and ageing^35^ (Fig. 3A&C). Low levels of SBS8 were observed in a subset of samples, in agreement with previous reports in PM^33,34^. Indel profiles were dominated by clock-like signatures including ID1 and ID2, also linked to replication slippage, with frequent contributions from ID5 and ID8. Less common indel signatures, including ID9 and ID10 of unknown aetiology, were detected in a subset of organoid–tumour pairs. Altogether, these results confirm that canonical mutational processes previously described in PM are consistently captured in organoid models.

Beyond these common processes, we identified distinct, sample-specific mutational signatures that were fully preserved in matched organoids. Notably, organoid model WTSI-MESO_002 and its paired tumour uniquely exhibited SBS2 and SBS13, associated with APOBEC cytidine deaminase activity (Fig. 3A). These signatures are characterised by enrichment of C>T and C>G mutations at TpCpX trinucleotides and have previously been reported in a subset of PMs^33,34^.

Similarly, organoid model WTSI-MESO_035 uniquely displayed the indel signature ID6, indicative of homologous recombination (HR) deficiency^35^, which was also present in the matched tumour (Fig. 3A). Consistent with this, both the organoid and tumour were classified as BRCA2-type HR-deficient and exhibited loss of RAD51B, a key mediator of RAD51-dependent repair (Fig. 2A; Supplementary Table 4; Methods).

We also identified a hypermutated case, WTSI-MESO_098, comprising both the primary tumour and derived organoid, with a markedly elevated mutational burden of 25.9 mutations per Mbp (∼14-fold higher than the cohort median) (Fig. 3A). These samples presented loss of *MLH1*, a core mismatch repair (MMR) gene (Fig. 2A). Correspondingly, both samples exhibited strong contributions from SBS44, a signature diagnostic of MMR deficiency and microsatellite instability (MSI), alongside increased SBS1, SBS5, and SBS41 activity^35^ (Fig. 3A). Indel profiles were dominated by ID1 and ID2, reflecting characteristic insertion-deletion events at repetitive sequences seen in MMR-deficient tumours (Fig. 3A). We compared the hypermutant organoid-tumour pair with a previously reported hypermutated PM case (TCGA-UD-AAC1)^31^. Despite distinct genetic drivers of MMR deficiency (*MLH1* loss versus *MSH2* mutation, respectively), mutational profiles were highly concordant, particularly for SNVs, suggesting convergent mutational processes across hypermutant PM cases (Fig. 3E; Extended Fig. E3). Clinically, the donor corresponding to WTSI-MESO_098 exhibited an unusually prolonged survival (∼1000 days from diagnosis) following first-line immune checkpoint inhibitor (ICI) therapy (nivolumab plus ipilimumab), substantially exceeding the survival observed in other cohort members (total cohort median=300 days, IQR=191-546, n=24 unique donors; cohort with ICI first line median=574 days, IQR=333-775, n=5 unique donors) (Supplementary Table 1). This observation is consistent with evidence from other tumour types linking MMR deficiency and hypermutation to improved responses to immune checkpoint inhibitors, likely mediated by an increased neoantigen burden^36,37^.

These data demonstrate that mesothelioma organoids faithfully recapitulate both common and rare mutational processes present in primary tumours, including age-related mutagenesis, APOBEC activity, homologous recombination deficiency, and mismatch repair deficiency. Importantly, organoids capture clinically relevant outlier phenotypes, supporting their utility as diverse representative models for studying tumour biology and therapeutic response in PM.

### Pleural mesothelioma organoids are enriched for cell-cell interaction pathways

We next compared the transcriptional profiles of PM organoids and primary PM cell lines. Across expressed protein-coding genes (n = 16,063), both models showed broadly similar profiles (mean pairwise Pearson r: organoid-organoid 0.86, cell line-cell line 0.88, organoid-cell line 0.85), consistent with their shared tumour origin despite deriving from independent cohorts of donors (Supplementary Fig. S9A). Restricting the analysis to the top 500 most variable genes lowered correlations (0.37, 0.49 and 0.25, respectively) and revealed a clearer separation between model systems, with cell lines appearing more homogeneous, whereas organoids retain greater inter-donor variability. Consistently, organoids and primary cell lines formed two distinct clusters, indicating that model type outweights subtype-associated variation (Supplementary Fig. S9A&B). While cell lines showed partial stratification consistent with epithelioid and non-epithelioid phenotypes, such separation was not evident in organoids, which may stem from the limited number of non-epithelioid organoids. Notably, organoids derived from the same donor but different anatomical sources (tumour versus pleural fluid, WTSI-MESO_006/009 and WTSI-MESO_046/047) clustered closely, suggesting that inter-patient variation predominates over derivation context.

Differential expression analysis identified 1,502 genes upregulated in organoids and 696 upregulated in primary cell lines, with limited overlap in enriched pathways (Fig. 4A&B; Supplementary Table 9&10). Cell lines preferentially upregulated gene sets associated with cell-substrate interactions, including focal adhesion, integrin signalling, and actin cytoskeleton organisation, consistent with adaptation to 2D culture^38^. These pathways included key regulators such as *THBS1* and *CNN2*, linked to extracellular matrix (ECM) engagement and adhesion (Fig. 4B&C). In contrast, organoids showed enrichment for cell-cell interaction and junctional signalling, including adherens junctions, tight junctions, and gap junction pathways, indicative of enhanced intercellular connectivity and tissue-like organisation. Representative genes included *PARD6B* and *CRB3*, involved in cell-cell junction formation, and polarity^39^. Organoids and primary cell lines appear to regulate ECM homeostasis differently, showing differential expression of genes involved in ECM formation and regulation (e.g., *COL14A1*, *SERPINE1*), suggesting active matrix remodelling in a 3D organoid context^40^ (Fig. 4B&C). Finally, shared pathways between models were limited, although Rap1 signalling (implicated in both cell-cell and cell-matrix adhesion^41^) was enriched in both systems.

**Figure 4.**
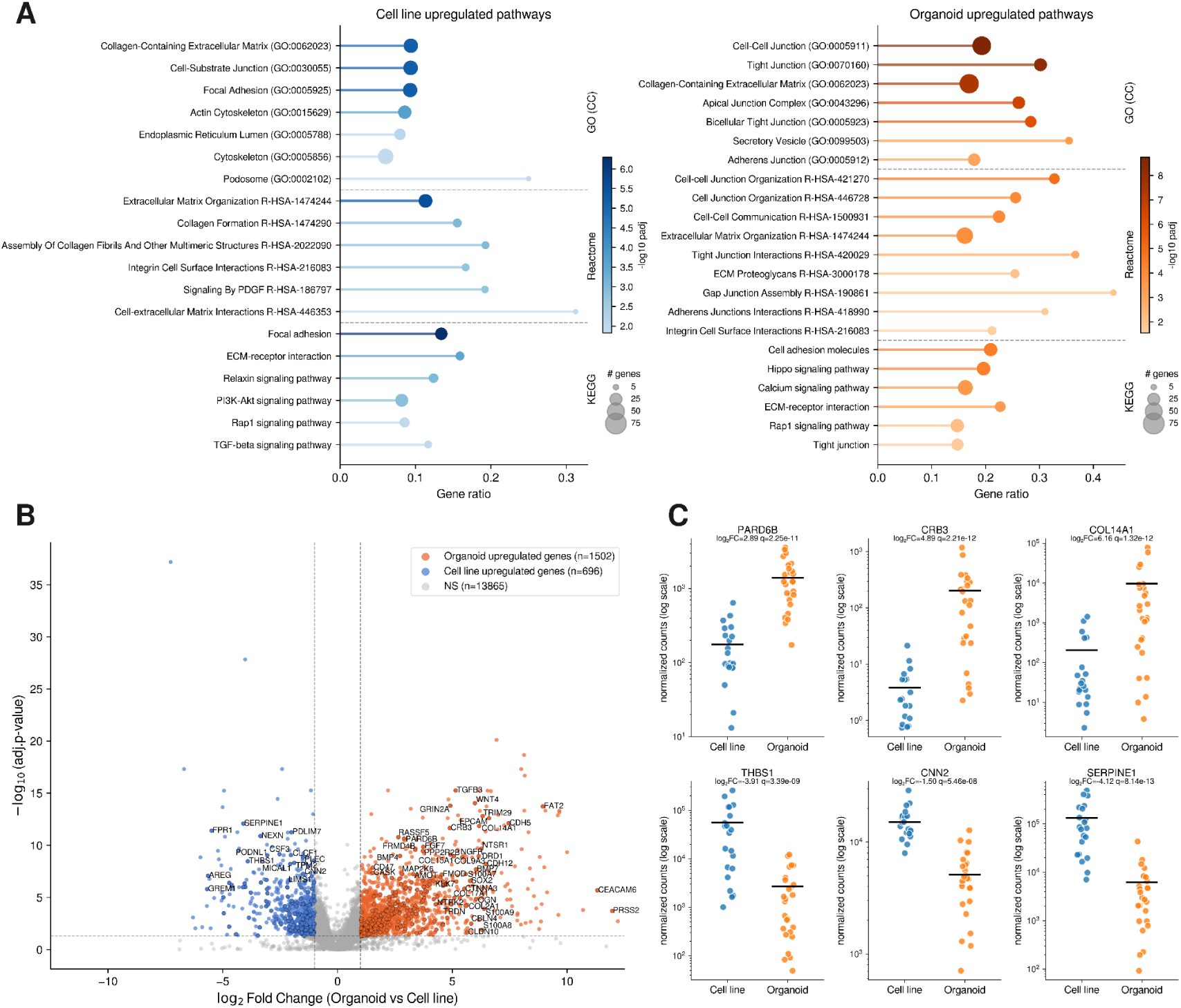
Transcriptomic analysis and differential gene expression across mesothelioma organoids and primary cell lines. (A) Over-representation analysis (ORA) of differentially expressed genes between organoids and primary cell lines across Gene Ontology (GO), Reactome, and KEGG pathways. Enriched gene sets are shown for genes upregulated in organoids (orange) and in primary cell lines (blue). The x-axis represents gene ratio (proportion of the term’s genes present), point size indicates the number of genes per pathway, and colour intensity reflects statistical significance (−log_10_ adjusted p-value). (B) Volcano plot of differential gene expression between organoids and primary cell lines. The x-axis shows log_2_ fold change and the y-axis −log_10_ adjusted p-value. Genes significantly upregulated in organoids (padj < 0.05, |log_2_FC| > 1) are shown in orange, and those upregulated in primary cell lines in blue; non-significant genes are shown in grey. Genes belonging to significantly enriched pathways are outlined in black, and selected genes with high fold change and significance are labelled. Dashed lines indicate significance thresholds (|log_2_FC| = 1 and adjusted p-value = 0.05). (C) Normalised expression of selected representative genes across sample groups. Each point represents an individual sample (n=26 organoids, n=21 cell lines), and horizontal bars indicate group medians. Colours denote model type (organoids, orange; cell lines, blue).

Overall, these results indicate that while organoids and primary cell lines share a common mesothelioma transcriptional backbone, they diverge substantially in pathways reflecting their structural context, with organoids favouring cell-cell interactions and tissue architecture, and cell lines exhibiting features consistent with 2D adhesion and substrate-dependent growth.

### Identification of pleural mesothelioma core fitness genes

To define the genetic dependencies required for pleural mesothelioma viability and growth, we performed genome-wide CRISPR-Cas9 dropout screens across 24 PM models comprising 15 organoids and 9 low-passage primary cell lines. Following stringent quality-control filtering, 23 models (14 organoids and 9 primary cell lines) were retained for downstream analyses (Fig. 5; Extended Fig. E4; Supplementary Fig. S10). The screens demonstrated excellent performance in discriminating reference essential and non-essential genes, with comparable results obtained in organoids and primary cell lines (mean AUROC 0.945, 0.973; mean AUPR 0.946, 0.975, respectively) (Supplementary Fig. S10), consistent with previously reported screens in tumour organoids and cell lines^11,42^. This demonstrates that high-quality genome-wide dependency maps can be generated using patient-derived PM organoids and PM primary cell lines.

**Figure 5.**
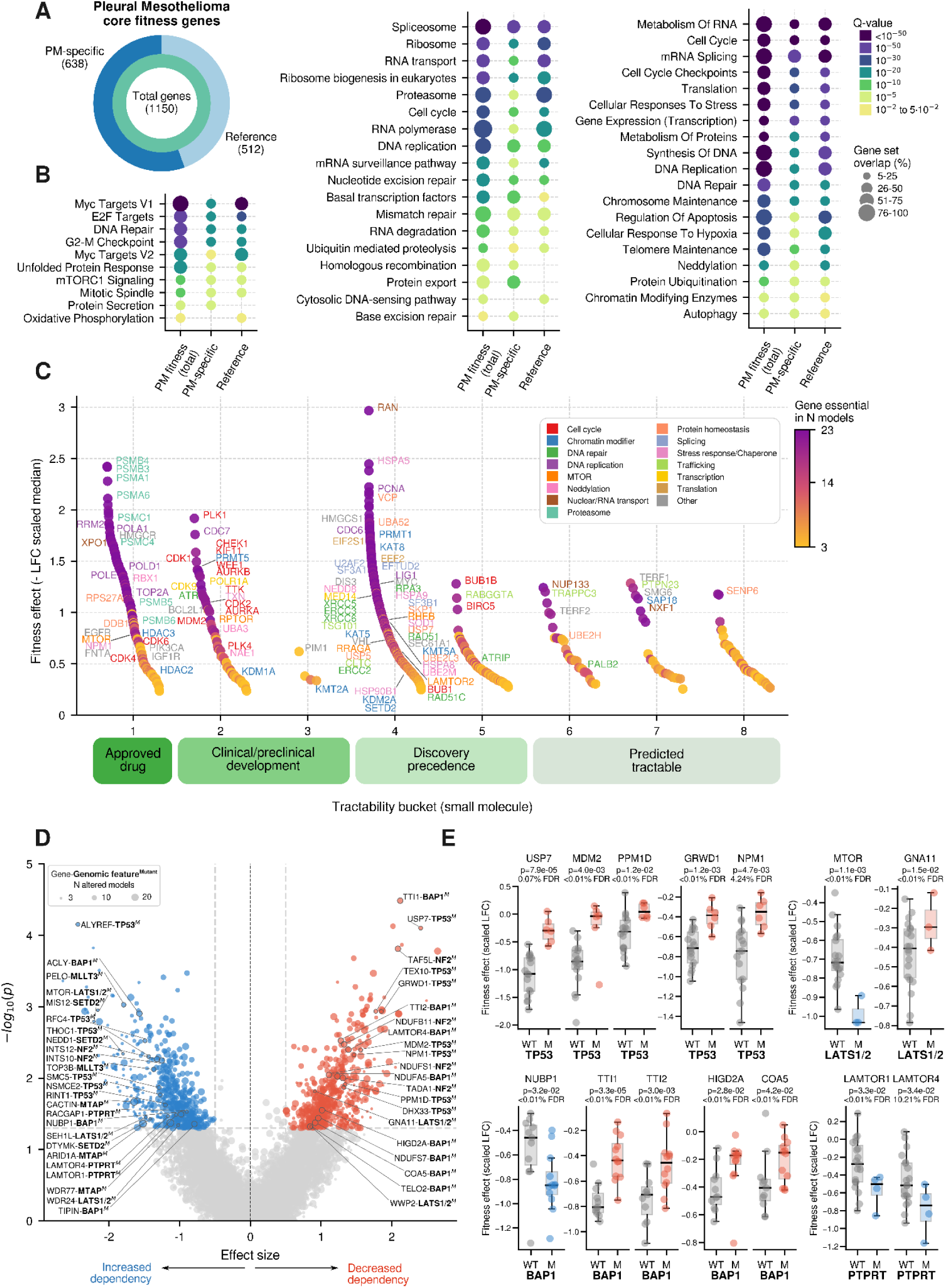
Prioritisation of pleural mesothelioma fitness genes and identification of therapeutic vulnerabilities. (A) Classification of pleural mesothelioma (PM) core fitness genes. Of the 1,150 PM core fitness genes, 638 were classified as PM-specific fitness genes and 512 as reference core fitness genes typically shared across multiple tumour types. (B) Functional enrichment analysis of total PM core fitness genes, as well as PM-specific and reference gene lists subclasses. Dot plots show relevant, significantly enriched biological processes grouped into major functional categories using MSigDB (left), KEGG (center), and Reactome (right) classifications. Dot size represents the gene set overlap, and colour indicates statistical significance (adjusted *P* value (Q-value)). (C) Target prioritisation and tractability assessment of PM fitness genes. Identified priority targets were classified according to small molecule tractability evidence and assigned to tractability buckets: approved therapeutic target (Bucket 1), targets with compounds in clinical or preclinical development (Buckets 2–3), targets with discovery precedence (Buckets 4–5), and targets predicted to be tractable based on structural, chemical or biological evidence (Buckets 6–8). Colour scale indicates the number of models in which the gene is essential, text colour indicates the broad pathway or functional group of the gene, y-axis indicates the median of the negative fitness effect across models where the gene was considered essential, higher y-axis indicates a stronger fitness effect. (D) Volcano plot showing context-specific dependency associations between genomic features and CRISPR-Cas9 dependency effects across PM models. Negative effect size associations indicate context-dependent increases in dependency (synthetic lethality/sensitivity), whereas positive associations indicate reduced dependency (synthetic resistance/viability). Dot size indicates the number of models with a specific genomic feature, labels are depicted as CRISPR target (left) and altered genomic feature in bold (right). (E) Representative biomarker-CRISPR target associations illustrating context-specific dependency responses. Plots show differential fitness effects of selected CRISPR targeted genes according to genomic status, between wild-type (WT) and altered (M, mutated or lost). Scaled log fold change (LFC) values are shown for individual models; boxes indicate the median and interquartile range (IQR), whiskers extend to 1.5 × IQR, and statistical significance from biomarker association analysis is detailed.

Across the screened models, we identified a median of 1,904 fitness genes in organoids (range 1,049–2,519 genes per model) and 2,366 fitness genes in primary cell lines (range 1,849–2,696 genes per model) (Extended Fig. E4A&B; Supplementary Table 11). To identify genes that were consistently required across PM models, we applied the Adaptive Daisy Model (ADaM)^10^, which determined that dependency in at least 17 models represented the optimal threshold for core fitness classification (Extended Fig. E4C). Using this approach, we identified 1,150 PM core fitness genes, among which 638 (55.5%) were classified as PM-specific (Fig. 5A; Supplementary Table 12). Such cancer-specific dependencies may represent more attractive therapeutic targets than broadly essential genes^10^.

Functional enrichment analysis revealed that PM core fitness genes were enriched in fundamental cellular processes required for tumour maintenance (Fig. 5B; Supplementary Table 13). Prominent functional categories included cell-cycle regulation, including DNA replication, cell-cycle checkpoints and chromosome maintenance; RNA splicing and mRNA processing; ribosome biogenesis and transcriptional regulation; DNA repair pathways, comprising nucleotide excision repair, base excision repair, mismatch repair and homologous recombination; and multiple proteostasis-related pathways, including translation, protein trafficking, ubiquitination, neddylation, proteasomal regulation and unfolded protein response signalling. These findings define the principal biological processes that underpin PM survival and proliferation. To identify candidate therapeutically actionable dependencies, we integrated CRISPR-Cas9 fitness data with target tractability scores for small-molecule, antibody and PROTAC modalities^43^. To broaden the scope for candidate target identification, genes classified as essential in at least three PM models were evaluated and prioritised according to dependency strength, prevalence and tractability criteria (Fig. 5C; Supplementary Fig. S11&S12; Supplementary Table 14). This analysis identified multiple priority targets with established therapeutic potential, including genes for which approved drugs already exist (e.g., *CDK4, CDK6, HMGCR, FNTA, MTOR, RRM2, TOP2A, PSMB5, PSMC1, RBX1, XPO1*) or for which compounds are currently in clinical (e.g., *PRMT5, MDM2, CDK2, CDC7, ATR, WEE1, PLK1, BCL2L1, CDK9, NAE1/UBA3, TXN*) or preclinical (e.g., *HMGCS1, RHEB, VCP, RAN, SOD1*) development. In addition, numerous PM dependencies were assigned to high-confidence tractability buckets. These include targets with clinically approved compounds, as well as targets that lack one but have compelling biological rationale and favourable tractability (Fig. 5C; Supplementary Fig. S11&S12; Supplementary Table 14). These prioritisation analyses were based primarily on dependencies recurrently observed across PM models and therefore focus on targets with potential relevance to broader patient populations. These results provide a framework for the rational development and repurposing of targeted therapies in pleural mesothelioma.

Whereas PM core fitness genes define vulnerabilities that are broadly shared across mesothelioma models, additional dependencies may arise only within particular genetic PM backgrounds. To identify these context-specific vulnerabilities, we performed systematic biomarker analyses testing for associations between CRISPR-Cas9 dependency profiles and cancer driver genomic alterations. Molecular features comprised SNVs, indels, CNAs and SVs in cancer driver or PM-relevant genes. Given the low number of genomic alterations in PM, individual alterations were rarely sufficiently recurrent to test in isolation, so we collapsed alterations per gene into two binary categories reflecting the expected direction of functional consequence (mutated/lost and gain) and merged paralogs or co-altered genes into composite features (see Methods). Genes were included when classified as essential or non-essential in at least three models, while genomic features were required to be present or absent in at least three models.

Genotype-specific dependency analysis recovered several coherent relationships across recurrent mesothelioma driver contexts. Associations with TP53 status provided internal validation of the screen. Loss of the p53 negative regulators MDM2, PPM1D and USP7 was selectively deleterious in TP53 wild-type models, but comparatively neutral in TP53-altered models^44,45^ (Fig. 5D&E; Supplementary Table 15). A similar pattern was observed for the loss of multiple ribosome-biogenesis factors, including *NPM1, GRWD1, DHX33* and *TEX10*, consistent with activation of p53-dependent nucleolar stress responses^46–49^. Conversely, TP53-altered models showed increased dependency on genes involved in DNA replication, fork progression and genome stability, including *SMC5, NSMCE2, RFC4, RINT1, ALYREF, and THOC1*, consistent with increased reliance on compensatory replication-stress and DNA-damage-response pathways when p53 function is impaired^50–52^.

Hippo pathway alterations also produced an interpretable pattern of dependencies. LATS1/2-altered models showed selective dependency on MTOR and on the GATOR2 subunits WDR24 and SEH1L, implicating amino-acid sensing and mTORC1 activation as vulnerabilities in this genetic context^53–57^ (Fig. 5D&E; Supplementary Table 15). In contrast, loss of WWP2 and GNA11 was better tolerated in LATS1/2-altered models than in wild-type models. Since both genes act upstream of LATS1/2 to promote YAP activity, their reduced impact in LATS1/2-altered cells is consistent with epistasis once the downstream Hippo kinase brake has already been lost^58,59^.

BAP1-altered models showed a distinct pattern, with selective dependencies involving metabolism and genome maintenance. Sensitivity associations included ACLY, consistent with altered glycolytic and lipid-biosynthetic requirements following BAP1 loss^60,61^ (Fig. 5D&E; Supplementary Table 15). Additional dependencies included TIPIN and NUBP1, implicating replication-fork protection and repair enzyme maturation, consistent with a DNA-repair-compromised phenotype^62–64^. Conversely, loss of genes involved in oxidative phosphorylation and mTORC1 activation was better tolerated in BAP1-altered models, including *NDUFA5, NDUFS7, COA5, HIGD2A, TTI1, TTI2, TELO2* and *LAMTOR4*, consistent with reduced dependence on oxidative phosphorylation and altered mTORC1 signalling following BAP1 loss^60,65,66^ (Supplementary Table 15). Other recurrent alterations in PM also showed genotype-specific dependencies, such as MTAP, MLLT3, NF2 and PTPRT. PTPRT-altered models, for example, were selectively dependent on the Ragulator components LAMTOR1 and LAMTOR4, consistent with the PI3K-AKT activation reported following PTPRT loss^67,68^ (Fig. 5D&E; Supplementary Table 15).

These associations identify genotype-specific therapeutic opportunities and provide mechanistic insight into the molecular determinants of PM fitness. In contrast to PM-specific dependencies, which are shared across a large proportion of mesothelioma models, these biomarker-associated dependencies represent vulnerabilities that are contingent upon specific molecular alterations and may therefore provide opportunities for patient stratification and precision cancer medicine.

### Pharmacological profiling identifies broad and genotype-specific therapeutic vulnerabilities in pleural mesothelioma

Finally, we sought to identify therapeutically actionable vulnerabilities in pleural mesothelioma using a pharmacological screening platform^9,69^. Drug responses were assessed across 36 models, comprising 16 organoids and 20 low-passage primary cell lines. A library of 106 compounds was assembled to cover pathways and biological processes relevant to cancer and PM biology, including both approved and investigational agents, as well as targeted therapies and conventional cytotoxic drugs (Fig. 6A; Supplementary Table 16). Drug sensitivity was quantified using dose-response assays from which half-maximal inhibitory concentration (IC_50_) and area under the dose-response curve (AUC) values were calculated. Compounds were subsequently grouped according to their primary target pathway to identify biological processes that represent common pharmacological vulnerabilities across PM models. Analysis at the pathway level demonstrated that several target classes consistently reduced viability across a substantial proportion of screened models (Fig. 6B). These included inhibitors of NEDDylation, YAP/TEAD signalling, chromatin regulation through histone deacetylase inhibition, protein stability and degradation pathways, IGF1R signalling, cytoskeletal regulation, PI3K-mTOR signalling, mitosis and cell-cycle progression. These findings indicate that PM exhibits recurrent dependence on pathways controlling proteostasis, cell-cycle regulation, growth signalling and transcriptional and epigenetic regulation. In contrast, platinum-based chemotherapy and pemetrexed, which historically formed the backbone of systemic therapy for PM, demonstrated comparatively limited activity across the screened models (Fig. 6B&C). This observation is consistent with the modest clinical efficacy of cytotoxic chemotherapy in this disease and highlights the potential value of alternative targeted therapeutic approaches^70^.

**Figure 6.**
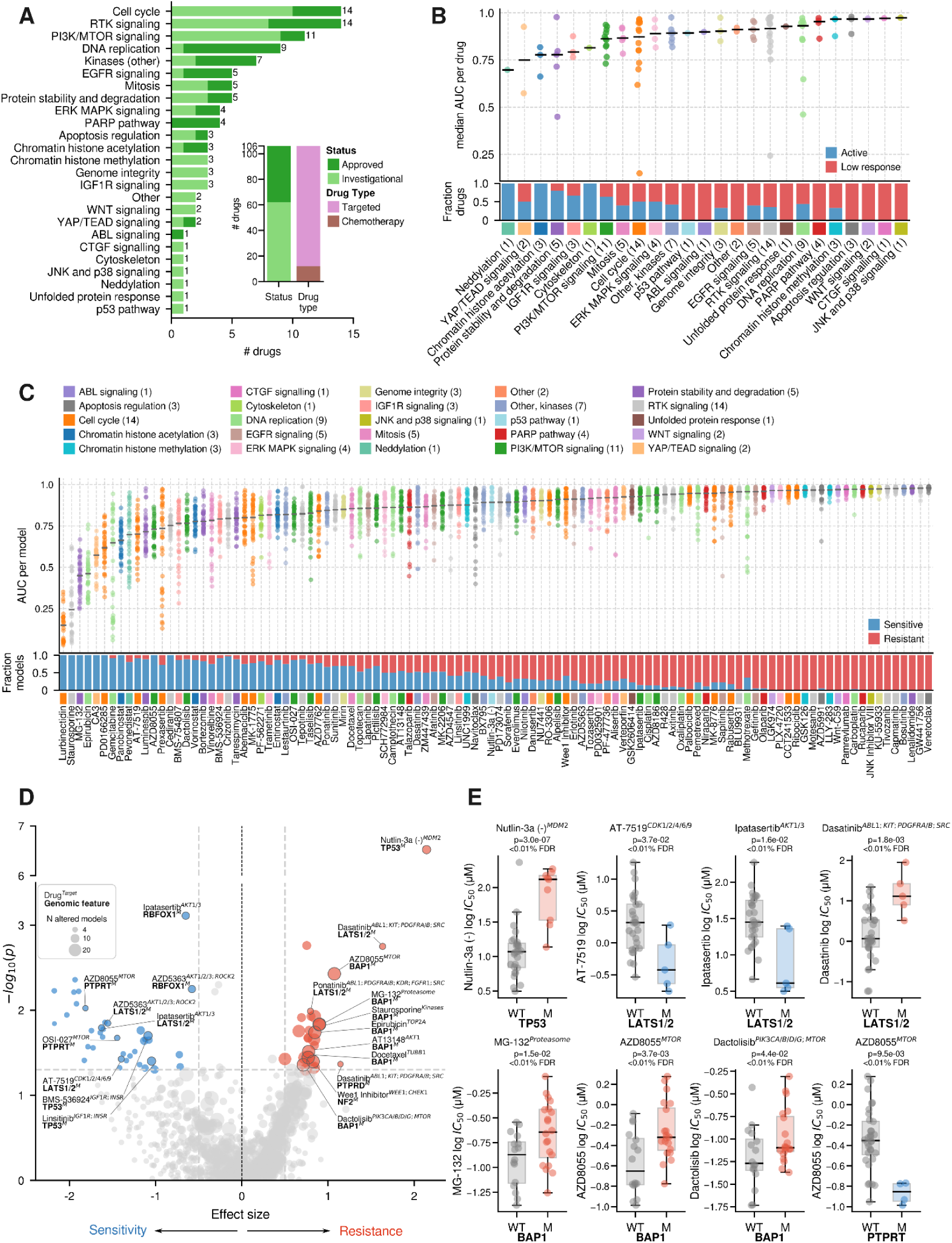
Pharmacological profiling identifies broad and context-specific therapeutic vulnerabilities in pleural mesothelioma. (A) Composition of the curated drug screening library. Compounds were grouped according to mechanism of action and biological pathway targeted. Light green indicates investigational compounds and dark green approved compounds. Pink denotes targeted agents and brown conventional cytotoxic chemotherapy drugs. (B) Pathway-level summary of drug responses across PM models. Upper panel shows the distribution of area under the dose-response curve (AUC) values for all compounds grouped by pathway. Individual drugs are represented by dots, and black bars indicate median pathway-level AUC values. Lower panel shows the proportion of drugs classified as active (blue) or low response (red) within each pathway category. (C) Drug activity across individual compounds. Upper panel displays AUC values for each screened drug across all models (n=36), while the lower panel shows the proportion of sensitive and resistant model responses observed for each compound. Black line indicates median AUC across all models for each drug. (D) Volcano plot showing biomarker-drug associations identified by pharmacogenomic analysis. Negative effect sizes indicate context-dependent sensitivity, whereas positive effect sizes indicate context-dependent resistance. Priority score thresholds are indicated and selected examples are labelled. Dot size indicates the number of models with a specific genomic feature, labels are depicted as indicated in the legend. (E) Representative biomarker-drug associations illustrating context-specific pharmacological responses. Plots show differential drug sensitivity according to genomic status, between wild-type (WT) and altered (M, mutated or lost). Log IC50 values are shown for individual models; boxes indicate the median and interquartile range (IQR), whiskers extend to 1.5 × IQR, and statistical significance from biomarker association analysis is detailed.

Several compounds showed particularly strong activity across the PM model panel (Fig. 6C and Supplementary Fig. S13). As expected, these included the broad-spectrum kinase inhibitor staurosporine and the proteasome inhibitor MG-132, both of which produced marked reductions in viability across most models, consistent with their well-established broad cytotoxic and antiproliferative effects in cancer cells^8^. Additional highly active compounds included the transcription inhibitor lurbinectedin, the DNA-damaging topoisomerase II inhibitor epirubicin and the YAP1 inhibitor CA3. Strong activity was also observed for the HDAC inhibitor panobinostat, the NEDDylation inhibitor pevonedistat targeting NAE1, the CDK inhibitor AT-7519, the HSP90 inhibitor luminespib, the mTOR inhibitor AZD8055, the WEE1 cell-cycle inhibitor PD0166285 and the pyrimidine antimetabolite gemcitabine. These findings identify multiple therapeutic strategies with potential relevance across PM subtypes.

Whereas several compounds demonstrated activity across a broad range of PM models, additional therapeutic opportunities may be restricted to specific genomic backgrounds. To identify such context-specific drug vulnerabilities, we performed systematic biomarker analyses testing for associations between drug sensitivity and recurrent cancer driver alterations. To focus on biologically relevant associations, drugs were included if classified as sensitive in at least three models, while genomic features were required to be present or absent in at least four models. Drug sensitivity was evaluated using IC_50_ values and tested against the presence or absence of recurrent genomic alterations (Fig. 6D&E; Supplementary Table 17).

The pharmacogenomic analysis recovered genotype-specific drug-response relationships. The clearest example involved TP53 status: TP53 wild-type models were selectively sensitive to the MDM2 inhibitor Nutlin-3a, whereas TP53-altered models were comparatively resistant (Fig. 6D&E; Supplementary Table 17), consistent with the selective MDM2 dependency identified in the CRISPR screen and with the requirement for functional p53 signalling downstream of MDM2 inhibition. This concordance provides an expected positive control for the biomarker analysis^71^.

Hippo pathway alteration was associated with differential response to several targeted agents. *LATS1/2*-altered models showed selective sensitivity to the multi-CDK inhibitor AT-7519, which is particularly potent against CDK9, consistent with reliance on transcriptional elongation downstream of constitutive YAP/TAZ-TEAD activity^72,73^ (Fig. 6D&E; Supplementary Table 17). *LATS1/2*-altered models were also selectively sensitive to the AKT inhibitors ipatasertib and capivasertib, implicating PI3K-AKT-mTOR signalling as a pharmacological vulnerability in this context and consistent with the MTOR dependency identified in the CRISPR screen^53–56^ (Fig. 6, 5D&E). Conversely, LATS1/2-altered models were less sensitive to dasatinib and ponatinib, both of which target SRC-family kinases among other targets, consistent with the loss of the LATS-dependent route through which SRC inhibition restrains YAP^74,75^ (Fig. 6D&E; Supplementary Table 17).

Drug associations with *BAP1* alteration were dominated by resistance rather than sensitivity. BAP1-altered models were less sensitive to several agents with distinct proximal targets but shared reliance on apoptotic execution, including staurosporine, epirubicin, docetaxel and MG-132, consistent with the reduced apoptotic competence reported following BAP1 loss^76,77^ (Fig. 6D&E; Supplementary Table 17). BAP1-altered models also showed reduced sensitivity to inhibitors of the PI3K-AKT-mTOR axis, including AZD8055, AT13148 and dactolisib, mirroring the better tolerance of loss of mTOR-related genes observed in the CRISPR screen^65,66^ (Fig. 6, 5D&E). Other recurrent alterations in PM also showed genotype-specific drug responses, such as NF2 and PTPRT. PTPRT-null models, for example, were more sensitive to the mTOR kinase inhibitors OSI-027 and AZD8055 (Fig. 6D&E; Supplementary Table 17), consistent with the mTORC1 dependency identified in the CRISPR screen^68^.

These findings demonstrate that pharmacological vulnerabilities in pleural mesothelioma arise both from dependencies broadly shared across tumours and from context-specific interactions associated with particular genomic alterations, providing a framework for both population-wide and biomarker-guided therapeutic strategies.

## DISCUSSION

The prognosis of pleural mesothelioma remains poor with few validated molecular targets^1^. A major obstacle to progress has been the scarcity of physiologically relevant experimental models that capture the biological diversity of the disease while remaining amenable to large-scale functional interrogation^13,14^. Owing to the difficulty of obtaining viable mesothelioma material, organoid cohorts reported to date have been small, largely derived from pleural effusions, and propagated using serum, autologous pleural fluid or conditioned medium retained from earlier culture^17,78,79^. To overcome these limitations, we undertook multi-centre recruitment and developed distinct derivation protocols for primary tumour tissue and pleural fluid together with a defined culture medium that supports long-term expansion without reliance on patient-derived supplements. Applying these new methods, we established the largest genomically annotated biobank of patient-derived mesothelioma organoids to date, validated against matched tumour and germline DNA and shown to retain the genomic alterations present in the originating tumours. We demonstrate the utility of this resource for integrated genomic, transcriptomic, genetic dependency and pharmacological profiling, identifying therapeutic vulnerabilities across mesothelioma and within molecularly defined subgroups. Our analyses provide a comprehensive map of mesothelioma vulnerabilities and identify candidate therapeutic strategies for both broad patient populations and molecularly defined subgroups.

We have shown that PM organoids faithfully preserve the key molecular features of their parental tumours. Across multiple orthogonal analyses, including driver gene alterations, copy-number architecture, mutational signatures, histopathology and lineage-marker expression, organoids showed remarkable concordance with matched tumours. Importantly, the organoid models established here preserved both common and rare genomic features of mesothelioma, ranging from canonical alterations in *BAP1*, *NF2* and *CDKN2A* to uncommon phenotypes such as homologous recombination deficiency, APOBEC activity and mismatch repair deficiency. The ability to model rare but potentially therapeutically relevant molecular states is particularly important in a cancer in which actionable genomic alterations are relatively uncommon.

Our analyses further demonstrate that organoids preserve biological complexity that appears diminished in established cell lines. Organoids preferentially expressed programmes associated with cell-cell interactions, polarity and tissue organisation, whereas cell lines displayed transcriptional features consistent with adaptation to two-dimensional growth and substrate attachment. These findings support the concept that culture context influences cellular transcriptional state and suggest that organoids may more accurately capture tumour-associated signalling networks^20^.

The generation of high-quality CRISPR-Cas9 dependency maps across patient-derived mesothelioma models represents another important advance. While large-scale cancer dependency resources have transformed the identification of therapeutic targets in common cancers, mesothelioma has remained substantially underrepresented in such datasets^10^. The dependency and tractability analyses converged on several classes of targets that appear particularly promising for therapeutic development. Notably, pathways involved in proteostasis and chromatin regulation emerged consistently across genetic and pharmacological screens. Although these findings merit further investigation, the concordance between orthogonal screening modalities increases confidence that these pathways represent genuine biological dependencies rather than artefacts of a particular experimental approach. Furthermore, the identification of numerous candidates within favourable tractability buckets indicates that a substantial fraction of mesothelioma dependencies may already be amenable to pharmacological intervention, either through drug repurposing strategies or through ongoing programmes in target discovery.

The pharmacological screen provides a systematic overview of therapeutic vulnerabilities across mesothelioma models, with broad activity among compounds targeting epigenetic regulation, YAP/TEAD signalling, proteostasis and PI3K-mTOR signalling, consistent with accumulating evidence implicating these pathways in mesothelioma pathogenesis^30,31,80,81^. Notably, platinum-pemetrexed chemotherapy showed relatively limited activity compared with multiple targeted approaches. Although direct comparisons between *ex vivo* systems and clinical outcomes should be interpreted cautiously, this is consistent with the modest benefit historically achieved with chemotherapy in PM and reinforces the need for mechanism-based strategies^1^. Several of the most active compounds converged on proteostasis, and the CRISPR screen identified multiple components of protein homeostasis networks as core fitness genes, providing orthogonal support for vulnerability. While previous studies suggested such a dependency^82–84^, interest in this area has waned. Our data suggest it is timely to revisit this biology, given the progress made in therapeutically targeting proteostasis over the last decade^85^.

Beyond identifying broadly shared vulnerabilities, CRISPR-Cas9 and pharmacological screens resolved differential responses in defined molecular contexts. Genetic and pharmacological perturbations are complementary, and in several instances provided orthogonal validation of the same underlying biology. The most informative associations were internally coherent, with functionally related genes or compounds moving together within a genotype, and converging on the established biology of recurrent mesothelioma drivers. The *TP53* associations illustrate this most clearly, confirming that canonical p53 biology remains detectable and pharmacologically exploitable in patient-derived models^45,71^. The LATS1/2 data suggest that loss of the Hippo brake redistributed dependencies, nominating mTOR-, AKT- and CDK9-directed agents for evaluation in Hippo-altered disease and SRC-directed agents for evaluation in tumours retaining LATS1/2^53–56,72,74^. BAP1 alteration produced a resistance-dominated phenotype, particularly to agents requiring mitochondrial apoptosis for efficient killing, consistent with the established non-nuclear role of BAP1 in stabilising IP3R3 and supporting ER-to-mitochondrial calcium transfer^76^, and presented reduced reliance on mTOR and oxidative phosphorylation, pointing to a metabolic and signalling rewiring. These associations nominate candidate stratification hypotheses, though they rest on modest model numbers and require prospective validation.

Several opportunities exist to extend this resource further in the future. Although this represents the largest functional mesothelioma organoid collections assembled to date, the number of models remains modest compared with pan-cancer resources. Continued expansion will increase power for biomarker discovery, particularly for rare genomic events. The cohort is also enriched for epithelioid tumours, with relatively few biphasic models and no sarcomatoid organoids. Whilst this likely reflects the limited availability of suitable sarcomatoid specimens, it remains possible that the culture conditions preferentially support the establishment or long-term propagation of epithelioid disease. Future work should therefore focus on extending representation of non-epithelioid mesothelioma and determining whether additional niche factors are required to support sarcomatoid growth *ex vivo*.

A further priority is to broaden the diversity of patients represented. Although recruitment was national, these models were derived within a single health system from a predominantly European-ancestry population, reflecting a broader imbalance in cancer model resources^86,87^. As genetic ancestry can influence drug response^88^, expansion to additional international centres will be important to establish the generalisability of the relationships described here. This is especially important given that much of the future global mesothelioma burden is expected to arise in countries with predominantly non-European populations, underscoring the need for models that better reflect the genetic diversity of affected patients. Future incorporation of co-culture systems, spatial profiling and in vivo validation will further extend the biological and translational insights that can be obtained from these models.

Standard cytotoxic chemotherapy will remain the principal treatment for most patients with mesothelioma worldwide owing to financial and logistical constraints on many healthcare systems, yet there are currently no validated biomarkers capable of identifying those most likely to benefit. By combining pharmacological profiling with matched genomic annotation, this resource provides a tractable platform for identifying predictive biomarkers of chemosensitivity alongside new therapeutic targets.

In summary, we have established a clinically annotated mesothelioma organoid biobank that faithfully captures the molecular diversity of pleural mesothelioma and supports large-scale functional interrogation. Through integrated genomic, transcriptomic, CRISPR and pharmacological analyses, we define core fitness genes, identify tractable therapeutic targets and uncover context-specific vulnerabilities associated with recurrent genomic alterations. Similar to other Cancer Dependency Map resources, these data provide a foundational resource for the mesothelioma research community and a framework for the development of both broadly applicable and biomarker-guided therapeutic strategies. More broadly, our study illustrates how patient-derived organoid platforms can bridge the gap between tumour genomics and functional precision oncology for rare and understudied cancers.

## Methods

### Sample acquisition

Patient tissue samples and associated clinical data used in this study were obtained from multiple Mesobank UK Tissue Collection Centres. Mesobank UK is a Research Tissue Bank that collects and distributes high-quality tissue and blood samples from patients with mesothelioma to support scientific research (www.mesobank.com). Mesobank’s participant recruitment procedures, participant information sheets, consent forms, and access policies - including requirements for processing research applications, conditions of access, and governance - have received a favourable opinion from the East of England - Cambridge Central Research Ethics Committee (REC reference: 18/EE/0161). Model derivation and analysis performed at the Wellcome Sanger Institute was supported by institutional ethics approval (IRAS ID:203519; REC 16/LO/1110).

### Pleural mesothelioma organoid derivation, expansion and culture

Methods to derive, culture and cryopreserve organoids from primary samples were optimised for PM from previous standard protocols^89–92^. Tissue samples were transferred from Falcon tubes with advanced DMEM and primocin to a glass Petri dish. Samples were washed 3 times with 10 mL of PBS. If the sample size allowed it, a small piece of tissue (0.5-1 cm) was sectioned and placed into a formalin pod to obtain primary tissue histology. A second small piece of tissue was obtained and further cut into smaller pieces (1-3 mm) for whole genome sequencing. The sample for sequencing was washed with 10 mL PBS, centrifuged at 800g for 2 minutes, the supernatant was aspirated, and the pellet was snap-frozen in dry ice for DNA extraction. To proceed with organoid derivation, 10 ml of digestion buffer were added to the main piece of tissue, which was then cut into very small pieces using a scalpel. Digestion buffer contains 0.5 ml collagenase (100mg/ml), 20 µl primocin (Invivogen (ant-pm-1)), 100 µl penicillin-streptomycin (Life Tech (15140122)), 10 µl Y-27632 (10mM) (Sigma Y0503) in 9.5 ml of Advanced DMEM/F-12 (Life Technologies (12634-028)). Sample and buffer were transferred to a 15 ml falcon tube, placed in a rotator/shaker at 37 ⁰C and digested for 1-4 hours, with periodic inspection under the microscope until sufficient digestion was observed. Then, using a 50 ml falcon tube, the mixture was filtered through a 1000 µm cell strainer to remove large, undigested fragments, and further filtered through a 100 µm cell strainer to remove smaller fragments. A pestle was used to aid filtration. Additionally, the initial Falcon was rinsed twice with 10 ml of PBS and passed through the cell filters. The digested sample was centrifuged at 800g for 2 minutes. The supernatant was aspirated, 30 mL of PBS were added, and the sample was centrifuged at 800g for 2 minutes. Optionally, if the pellet was very red, the pellet was resuspended in 5 mL of red blood lysis buffer (Thermo Fisher, 00-4333-57) and incubated for 5 minutes at room temperature, then 25 ml of PBS were added to dilute it, and an additional centrifugation step was repeated. After aspirating the supernatant, 2.5 ml of PBS were used to resuspend the pellet and transfer it to a 5 ml Eppendorf tube. This step was repeated, the sample in 5 ml of PBS was centrifuged at 800g for 2 minutes, and the supernatant was aspirated. If samples had an unconfirmed PM diagnosis, the pellet was resuspended in 1 ml of Recovery Cell Culture Freezing media (Thermo Fisher catalogue #12648010), placed in a CoolCell at −80 °C, and transferred to liquid nitrogen to continue with the protocol after a confirmed diagnosis. If samples had a PM confirmed diagnosis, the pellet was resuspended in cold 80% BME (80:20 in PM organoid media), seeding 200 µl per well of pre-warmed 6-well plate, with 10-15 µl droplets, in 1-5 wells of 6-well plates, depending on the size of the pellet. 6-well plates were incubated upside down for 15-30 minutes at 37 °C, and 2 ml of PM P0 media (PM organoid media supplemented with 2 µl/ml primocin, 10 µl/ml penicillin - streptomycin, 1 µl/ml Y-27632) per well was added. Media was changed twice a week with PM P0 media until the first passage.

Pleural fluid samples were filtered through a 1000 µm cell strainer to remove clots. Stripettes and pipettes used to manipulate pleural fluid were pre-wet 0.1% PBS/BSA to reduce sample loss. Filtered pleural fluid was centrifuged at 800g for 2 minutes in a conical-bottom bottle or a 50 ml falcon tube, the supernatant was aspirated and the pellet was resuspended in 40 ml PBS and transferred to a 50 ml falcon tube. The sample was centrifuged at 800g for 2 minutes. If the pellet was very red, the pellet was resuspended in 5 ml of red blood lysis buffer (Thermo Fisher, 00-4333-57) and incubated for 5 minutes at room temperature, then 25 ml of PBS were added to dilute it, and an additional centrifugation step was repeated. If the cell pellet was large, it was resuspended in 5-10 ml of advanced DMEM and part of the sample (10-20%) was transferred into a 15 ml falcon tube, PBS was added to obtain 10 ml, and the sample was centrifuged at 800g for 2 minutes. The supernatant was aspirated and the cell pellet was snap frozen in dry ice for DNA extraction. The remaining portion of the resuspended sample was centrifuged again and the supernatant was aspirated. A 100 µm cell stainer was pre-coated using 5 ml of 0.1% BSA/PBS to reduce sample loss. The pellet was resuspended in 5 ml of advanced DMEM and was filtered through a 100 µm cell stainer. Additionally, the tube was rinsed with 5 mL of advanced DMEM, and the rinse was filtered. The sample was centrifuged at 800g for 2 minutes and the supernatant was aspirated.

If samples had an unconfirmed PM diagnosis, the pellet was resuspended in 1 mL of Recovery Cell Culture Freezing media (Thermo Fisher catalogue #12648010), placed in a CoolCell at −80 °C, and transferred to liquid nitrogen to continue with the protocol after a confirmed diagnosis. If samples had a PM-confirmed diagnosis, the pellet was resuspended in cold 80% BME (80:20 in PM organoid media), seeding 200 µl per well of a pre-warmed 6-well plate, with 10-15 µl droplets, in 1-5 wells of 6-well plates, depending on the size of the pellet. 6-well plates were incubated upside down for 15-30 minutes at 37 °C, and 2 mL of PM P0 media (PM organoid media supplemented with 2 µl/ml primocin, 10 µl/ml penicillin - streptomycin, 1 µl/ml Y-27632) per well was added. Media was changed twice a week with PM P0 media until the first passage.

To be considered successfully derived, a model had to expand to more than 25 million cells and yield a bank of 25 cryovials; survive cryopreservation, re-expansion, and freeze-thaw cycles; remain free of contamination and fibroblast overgrowth; and pass model-tissue-blood concordance testing by immunohistochemistry and whole-genome sequencing. Genomic stability in culture was assessed by comparing pellets collected from eight patient-derived organoid models at two time points, separated by 4-11 passages. Moreover, organoids and their matched normal samples were verified using a 95 single-nucleotide polymorphism (SNPs) panel run on the Fluidigm 96.96 Dynamic Array IFC. Each organoid had to match its own normal sample (SNP Score ≥75%) and not match other samples.

Organoids were cultured in PM organoid media which contained: 95 ml organoid base media [500 ml Advanced DMEM (Life technologies, 12634-028), 5 ml 1M HEPES (Life technologies, 15630056, final conc. 10 mM), 5 ml 100X Glutamax (Life technologies, 35050038, final conc. 1X), 5 ml 100X CTS N-2 (Life Technologies A1370701, final conc. 1X)], supplemented with 2 ml RSPO3-FC CM (U-protein Express R001, avsbio, final conc. 2%), 1 ml Noggin CM (U-Protein Express BV N002, avsbio, final conc. 1%), 10 µl 5mM A83-01 (TOCRIS 2939, final conc. 500 nM), 1.67 µl 30mM SB202190 (Stem Cell Technologies 72634, final conc. 500 nM), 20 µl FGF2 100µg/ml in 0.1% BSA (Peprotech 100-18B, final conc. 20 ng/ml), 25 µl FGF7(KGF) 100µg/ml in 0.1% BSA (Peprotech 100-19, final conc. 25 ng/ml), 20 µl FGF10 500µg/ml (Peprotech 100-26-1000, final conc. 100 ng/ml), 10 µl recombinant human EGF 500µg/ml (Peprotech AF-100-15-1000, final conc. 50 ng/ml), 2 ml B-27 supplement 50X (Gibco 17504-044, final conc. 1X), 0.25 ml N-Acetylcysteine 500mM, (Sigma A9165-5g, final conc. 1.25 mM), 0.5 ml nicotinamide 1M (Sigma N0636-100g, final conc. 5mM).

To passage and expand the organoids, two previously described culture methods were employed: culture in 80% BME^90^ or suspension culture with 5% BME^69^. Media were aspirated, and 1 mL of TrypLE (Thermo Fisher, #12604013) was added per well of a 6-well plate. BME droplets were detached using a pipette tip or cell scraper, transferred to 5 mL tubes with P1000 pipette, and an additional 1 mL of TrypLE was used to rinse the well. The sample was incubated at 37 °C with periodic inspection under the microscope until the organoids were dissociated into small clumps of cells. The sample was centrifuged at 800g for 2 minutes, and the pellet was resuspended in cold 80% BME (Cultrex Sigma, 3533-010-02)(80:20 in PM organoid media) and seeded at 200 µl per well of a pre-warmed 6-well plate, with 10-15 µl droplets, in 2-6 wells of 6-well plates, depending on the size of the pellet. Alternatively, the pellets from 4-6 wells were resuspended in 20 mL of 5% BME in media and cultured in T75 ultra-low attachment flasks. Continuing with the 6-well plate method, the plate was incubated upside down for 15-30 minutes at 37 °C to allow BME to set, and 2 ml of PM organoid media was added per well. Plates were incubated at 37 °C, 5% CO2 and the media were changed twice a week with fresh PM organoid media until ready to passage.

### Primary PM cell line culture

Primary PM cell lines were obtained from Mesobank UK^28^ and cultured in RPMI-1640 (Sigma-Aldrich, R8758) supplemented with 10% FBS (Sigma-Aldrich, F7524), 100 U/ml penicillin, 100 μg/ml streptomycin (Sigma-Aldrich, P0781), 2 ng/ml hEGF (Gibco, PHG0315), 1 μg/ml hydrocortisone (Sigma-Aldrich, H4001) and 2 μg/ml heparin (Sigma-Aldrich, 2106), at 37 °C and 5% CO₂.

### Histopathological analyses

Immunohistochemical staining and haematoxylin and eosin staining were performed on the organoid models and, where available, the matched parent primary tumour tissues. For primary tissues, small pieces of tissue were fixed in 10% neutral buffer formalin solution (Sigma-Aldrich, HT501128) and subsequently embedded in paraffin. For organoid samples in culture, media was aspirated, and BME blobs were detached with 2 mL of cold Cell Recovery Solution (Corning, 354253) per well of a 6-well plate and were gently pipetted up and down, and subsequently incubated in the fridge for up to 1 hour with additional mild pipetting every 15 minutes. Samples were centrifuged at 800 g for 2 minutes, and the pellet was resuspended in 5 ml Sed-Fix working solution (Leica Biosystems, 3802954) and incubated at room temperature for 2 hours. Samples were centrifuged at 3000 rpm for 10 minutes; the supernatant was completely aspirated, and the pellet was resuspended in 500 µl of plasma (Sigma-Aldrich, P9523), 500 µl of thrombin (Sysmex, B4233-25) very gently to create a clot, and 5 ml of formalin solution and subsequently embedded in paraffin. Sections were stained with Leica Bond systems protocol F using Mesothelin (Leica, PA0373), Calretinin (Leica, PA0346), Wilms tumour protein WT1 (Leica, PA0562), Cytokeratin 5/6 (CK5/6) (Agilent, M723701-2), Podoplanin (D2-40) (Agilent, M361901-2) or with Ventana system using CK5/6 (Roche, 790-4554), WT-1 (Roche, 760-4397), Calretinin (Roche, 790-4467). Images were acquired using Leica Aperio AT2 scanning at 40x magnification, and representative images were obtained at 20x for primary tissues and 30x for organoids.

### Whole genome sequencing and analysis

#### Sequencing, alignment, variant calling and post-processing

DNA extracted from snap-frozen tumour tissues, blood, organoid pellets or cell line pellets were prepared for whole genome sequencing (WGS). Whole-genome paired-end sequencing reads (150bp) were generated using the Illumina HiSeq X Ten platform with an average coverage of 40x. Reads were aligned to GRCh38 human reference genome using the Burrows-Wheeler Alignment (BWA-MEM) tool^93^. PCR duplicates, unmapped and non-uniquely mapped reads were flagged before downstream analysis.

Single nucleotide variants (SNV) and insertions and deletions (INDELS) were identified using cgpCaVEManWrapper^94^ and cgpPindel^95^, respectively. AscatNgs^96^ output values were used as input to copy number and normal contamination for cgpCaVEManWrapper when run with matched normal blood.

Sample-specific germline variants were filtered using matched blood samples where available. For samples lacking matched blood, an in silico normal sample generated from the GRCh38 human reference genome was used instead. In those samples, variants in gnomAD v3.1.1 and 1000 Genomes with population-level AF (Allele Fraction) >0.001 were used to filter germline variants. For all samples, additional germline variants and technology-specific artefacts were removed by filtering against a panel of 100 unrelated normal samples (ftp://ftp.sanger.ac.uk/pub/cancer/dockstore/human/GRCh38_hla_decoy_ebv/SNV_INDEL_ref_GRCh38_hla_decoy_ebv-fragment.tar.gz).

Post-processing filters were applied using the in-house tool cgpCaVEManPostProcessing (github.com/cancerit). Unbiased analysis of mutant and wild-type reads found at the loci of the SNVs and Indels were assessed within a group of related samples using vafCorrect^97^. Only mutations that passed quality control criteria, overlapping nuclear chromosomes, did not overlap blacklisted regions, and were not flagged as overlapping the normal panel germline (NPGL) were kept. For samples lacking matched blood, mutations were included for analysis if their VAF (Variant Allele Fraction) was greater than 0.15.

Blacklisted regions for GRCh38 reference genome, including the ENCODE Blacklist^98^ and GRC Exclusions, were obtained from UCSC Genome Browser at genome.ucsc.edu.

#### Mutation burden

To compute the mutation burden (mutations per Mbp) for small variants, we divided the number of variants per sample by the total number of known positions (non-N) in the GRCh38 reference genome based on ncbi.nlm.nih.gov/grc/human/data after removing blacklisted positions.

#### Identification of driver genes and driver single-nucleotide and indel variants

Cancer driver genes were obtained from the Cancer Driver Gene list from Cell Model Passports (CMP)^99–101^. Mutations were annotated as driver variants if they overlapped the genomic coordinates of a mutation with the same effect (e.g. nonsense, missense) in the Cancer Driver Mutations library from CMP. For genes with Loss of Function (LoF) or Ambiguous method of action in the Cancer Driver Gene list, mutations were annotated as driver variants if the mutation had a LoF effect (e.g. stop gain, stop loss, frameshift). Mutations in mesothelioma-associated (non-CMP driver) genes of interest (CDKN2B and MTAP) were flagged if the mutation had a LoF effect. Mutations were annotated as cancer predisposition variants if they overlapped the genomic coordinates of a mutation with the same effect (e.g. nonsense, missense) in the Cancer Predisposition Variants (CPV) list from CMP^102^. Similarly, for genes with LoF or Ambiguous method of action in the CPV list, mutations were annotated as CPV if the mutation had a loss-of-function effect (e.g. stop gain, stop loss, frameshift).

For illustrative purposes of heatmap driver alterations, driver genes were selected based on the number of mutated organoids with matched blood containing point mutations, structural variants and copy number high-loss or amplification alterations. Those genes containing 3 or more alterations, together with additional genes of interest, were displayed, sorted by total alteration count across the organoid cohort.

#### Copy number and genomic instability analysis

Gene-level copy number alterations (CNAs), segment-level copy number alterations (SCNAs), purity, ploidy, structural variants, microsatellite instability status and homologous recombination deficiency status were obtained for each sample independently using Hartwig Medical Foundation tools, available on github.com/hartwigmedical/hmftools. Gene-level CNA, SCNA, purity, ploidy and microsatellite instability status were obtained from Purple (≥v2.54)^103^, using matched blood as germline where available. Structural variants in cancer driver genes were computed with Linx (v2.1)^104^ using GRCh38 reference genome. All other parameters were kept as default. Disruptions and homozygous deletion disruptions were considered drivers if driver likelihood was 1, as computed by Linx. Homologous recombination deficiency status was predicted using Chord (v2.0.3)^105^.

#### Gene-level copy number analysis

For each sample, gene-level copy number (CN) was categorised into five different groups (CNclass) according to the following formula^11,101^: where CN is the minimum gene copy number, and ploidy is the ploidy of the sample. Rounded CN class values were classified as: 0 high loss, 1 loss, 2 neutral, 3 soft gain, 4 amplification. Genes with a fraction larger than 0.1 of their canonical transcript exonic sequence overlapping blacklisted regions were excluded. Canonical transcript identifiers and exonic coordinates were downloaded from GRCh38 Ensembl version 114 and 112, respectively. Copy number altered genes were annotated as drivers if the genes were found in the driver gene list or in MTAP and CDKN2B.

#### Segment-level copy number analysis

In order to compare segment-level copy number (SCNA) across the genome between samples, we intersected SCNAs with non-overlapping 100 Kbp genomic bins obtained from GRCh38 reference genome (bgreference v0.7; pybedtools v0.10.0). Bins with sequences containing 1% or more unknown nucleotides and/or bins overlapping blacklisted regions were excluded from further analysis. For each bin, the weighted CN mean was computed as the average of CN across CN segments in the bin normalised by the bin length. Ploidy-normalised CN values across autosomic bins were compared between organoid and matched tissue or early passage organoid using Pearson correlation (scipy v1.9.3).

#### Clonal fraction

For each SNV and Indel mutation, cancer cell fraction (CCF) (proportion of tumour cells in a sample that carry a specific mutation) was obtained using mutation VAF, sample total copy-number and purity, and the minor and major allele copy-number values obtained from the segmented copy-number data in the mutated position, according to the formulas detailed in Steele et al. (2019)^106^ and Dentro et al. (2021)^107^. Mutations with CCF > 0.8 were considered to be present in all cancer cells and classified as clonal. For each sample, the clonal fraction was computed as the proportion of clonal mutations.

#### LOH and biallelic mutations

Genes classified as tumour suppressor or ambiguous genes in the cancer driver list were annotated as containing loss of heterozygosity (LOH) if the gene showed the loss of one copy (CN classified as “loss”) and carried a loss of function mutation (e.g. stop gain, stop loss, frameshift) with variant allele frequency VAF ≥ 0.8 in organoids and cell lines or VAF ≥ 0.5 in tissues to account for differences in purity. Biallelic inactivated tumour suppressor or ambiguous genes were considered as those containing complete gene loss (CN classified as “high loss”), biallelic structural variants, or a loss of function mutation with allele frequency VAF ≥ 0.8 in organoids or VAF ≥ 0.5 in tissues.

### Public datasets analysis

Open variant calls from The Cancer Genome Atlas (TCGA) pleural mesothelioma cohort were downloaded from Genomic Data Commons at portal.gdc.cancer.gov. Substitutions and indels were obtained from WXS masked somatic mutations (GRCh38.p0). Gene and segment copy number data were obtained from ASCAT3 gene-level copy number files and copy number segments, respectively. Samples with available open access data and previously analysed by Hmeljak et al.^31^ were kept (n=70). Substitutions and indels were classified as drivers as previously described. CN gene categories were computed using CN gene data and purity estimates from CN segment data, and classified as previously described.

Mutation burden for established available pleural mesothelioma cell lines were obtained from CMP at cellmodelpassports.sanger.ac.uk. The following cell lines were included (n=25): IST-MES-2, NCI-H2373, NCI-H2595, NCI-H2722, NCI-H2731, NCI-H2369, NCI-H2461, NCI-H2591, NCI-H2795, MPP-89, NCI-H226, NCI-H2810, NCI-H2804, NCI-H2818, NCI-H2869, NCI-H290, MSTO-211H, NCI-H2452, NCI-H2803, ACC-MESO-1, JL-1, NCI-H28, RS-5, IST-MES1, NCI-H2052.

### Mutational signature analyses

De novo mutational signature extraction was carried out using sigprofiler tools suite. First, SNV and indel mutational profiles were obtained for samples with matched blood using SigProfilerMatrixGenerator (v1.2.31)^108^. SBS96 and ID83 mutational signatures were extracted using SigProfilerExtractor (v1.1.25)^109^ with GRCh38 as the reference genome and 1,000 NMF replicates. We tested solutions between 1 and a maximum of 20 and 15 signatures for SBS96 and ID83, respectively. All other parameters were used as default. The best manually curated solution was selected after decomposing all considerable solutions from SigProfilerExtractor into COSMIC v3.4 SBS96 and ID83 mutational signatures^110^ using SigProfilerAssignment (v0.1.9)^111^. Mutational profiles were compared using cosine similarity (1 - cosine distance)(scipy v1.12.0). To compare mutational profiles from TCGA tumours (whole exome) with our dataset (whole genome), we computed the profiles for both datasets, restricting mutation calls in exonic regions using SigProfilerMatrixGenerator (exome=True).

### RNA sequencing and analysis

Paired-end RNA-sequencing reads (75 bp) were quality-filtered and aligned to the GRCh38 reference genome using STAR (v2.5.0c)^112^ with default parameters (github.com/cancerit/cgpRna). Gene-level read counts were subsequently obtained from the resulting BAM files using RSEM (v1.3.3)^113^. Protein-coding genes were subsetted from raw count matrices, and genes with fewer than 10 counts in fewer than 3 samples were removed. Principal component analysis and sample-sample Pearson correlations were obtained on variance-stabilising transformation-normalised expression values across the 500 most variable genes with PyDESeq2 (v0.5.4)^114^. Differential expression analysis between model types, correcting for histological subtype, was performed on PyDESeq2. Multiple test correction using Benjamini-Hochberg and log2 fold change shrinkage was applied to obtain the differentially expressed genes with adjusted p-value < 0.05 and |log2 fold change| > 1. Over-representation analysis (ORA) was performed separately on upregulated and downregulated gene sets using gseapy (v1.1.13)^115^ against GO Cellular Component^116^, KEGG^117^, and Reactome^118^.

### CRISPR-Cas9 screen

#### Screening

We analysed a total of 15 organoid and 9 cell line models for CRISPR-Cas9 screens. All models were analysed in triplicate, except WTSI-MESO_007, which was analysed in duplicate. Detailed protocols for generating Cas9-expressing organoid cultures and for CRISPR-Cas9 library transduction in organoids, including process diagrams and example data, are available on protocols.io^119,120^ and were followed here. Stable Cas9-expressing cancer organoid lines were generated by transduction with lentiCas9-Blast (Addgene 52962), with polybrene (10 μg/ml) added to enhance transduction efficiency. Following overnight incubation, the lentivirus-containing medium was replaced with fresh complete PM organoid medium supplemented with Y-27632 (2.5 µM final concentration). Continuous blasticidin selection (Invivogen, ant-bl-1) was applied until the required number of cells for the next steps, including Cas9 activity assessment, had been reached. Cas9 activity was assessed using a fluorescent reporter assay (Addgene 67982 and 67981)^121^, and only organoid lines exhibiting greater than 75% Cas9 activity were carried forward for sgRNA library transduction. Screens used MinLibCas9 (Addgene 164896)^122^, a genome-wide human CRISPR-Cas9 library. It targets 18,761 genes using 2 optimal sgRNAs per gene and has a total of 37,522 gene-targeting and 200 non-targeting sgRNAs. The lentiviral volume required to achieve an MOI of ∼0.3 for the sgRNA library was determined by titration. The percentage of cells transduced was assessed using flow cytometry for BFP-positive cells. A total of 3.75 × 10^7^ Cas9-expressing organoid cells were transduced in triplicate using the lentiviral-packaged whole-genome sgRNA library (∼100× coverage) in medium supplemented with polybrene (10 μg/ml) and Y-27632 (2.5 μM). Following overnight incubation, cells were plated in fresh medium as a 5% BME2 suspension. Transduction efficiency (target 30%) was assessed by flow cytometry. Screens were then maintained under puromycin selection for ∼16 days (∼21 days total screen). A final selection efficiency of at least 60% by flow cytometry was required for the screen to pass. At the end of the screen, approximately 2.5 × 10^7^ cells per replicate were collected, pelleted, and stored at −80°C for downstream DNA extraction and sequencing.

Detailed protocols for generating Cas9-expressing cancer cell lines and for genome-wide CRISPR-Cas9 screening, including process diagrams and example data, are available on protocols.io^123–128^ and were followed here, adapted for use with the same MinLibCas9 library. These protocols implement the screening pipeline described by Behan et al.^10^. All main steps were equivalent to the organoid procedures described above, differing only in culture format and in minor parameters detailed in the referenced protocols: blasticidin titration, Cas9 transduction and activity assessment against the same >75% threshold, puromycin and sgRNA library titration, library transduction in technical triplicate at ∼100× coverage, puromycin selection requiring a final selection efficiency of at least 80% by flow cytometry, and subsequent expansion, sample collection and downstream processing.

#### Quality-control and data processing

To evaluate the quality of the screens, we first applied an sgRNA replicate quality control check following the protocols detailed in Behan et al.^10,11^. Using the 37,522 gene-targeting sgRNAs, we computed the pairwise Euclidean distance between sgRNA pairs for each gene. We then selected the top 2.5% genes (469 genes) whose sgRNAs were most consistent across replicates and models, independently for the organoid and cell line cohorts. Next, for each model, we used the gene-level fold-change values of these selected genes to compute all possible pairwise Euclidean distances between all available technical replicates. Replicates were highly consistent for all models (mean Euclidean distance range 6.1-23.7) tested except WTSI-MESO_009 (mean Euclidean distance 121.6), which was therefore excluded from downstream analyses.

The analysis of 14 organoid and 9 cell line models was further processed using CRISPRcleanR (github.com/francescojm/CRISPRcleanR) to normalise sgRNA counts and correct for gene-independent responses to CRISPR-Cas9 targeting, such as copy-number bias, and to obtain corrected logFC values^129^. Screen quality was assessed by ROC and precision-recall area under the curves to assess the ability to identify reference essential/non-essential genes. We used curated reference gene sets of essential and non-essential genes from BAGEL2^130^ adapted to our assay to match minLib library. Corrected values were then processed with BAGEL2 (github.com/hart-lab/bagel) to identify genes showing significant depletion in each model, and genes classified as essential in a given model were reclassified as non-essential if the gene was subject to a High Loss event in that model. Cancer driver genes were removed from the reference gene sets from the BAGEL2 steps onwards.

To improve comparability across models, gene-level LFCs were scaled using reference essential and non-essential genes as controls: for each gene, the median non-essential LFC was subtracted, and the result divided by the difference between the median non-essential and median essential LFCs.

We used the Adaptive Daisy Model (ADaM2) software^10^ to identify core fitness genes in pleural mesothelioma, using all models and their per-model essential gene classifications as inputs, defining the threshold at the crossover point. In total, 1,150 PM core fitness genes were identified, each being essential in at least 17 models. These genes were further classified according to whether or not they were present in the reference list of essential genes. Over-representation analysis (ORA) was performed on all PM core fitness genes, and the subsets of PM-specific or reference sets, using gseapy (v1.1.13)^115^ against MSigDB^131^, KEGG^117^, and Reactome^118^.

#### Tractability assessment

To prioritise gene candidates from the CRISPR-Cas9 screen as potential targets, we used tractability scores from Open Targets platform (ftp.ebi.ac.uk/pub/databases/opentargets/platform/latest/input/target/tractability/ version 2025-08-18). All unique genes with available tractability data identified as fitness genes in one model or more models were processed, and genes without available tractability data were discarded. Tractability data was classified into three main categories: small molecules (SM), antibodies (Ab), and PROTACs. For each category, genes were classified in a tractability bucket ranking from 1 (highest) to 10 (lowest) tractability score, and further classified into 3 groups (clinical precedence, discovery precedence and predicted tractable, or the equivalent for SM, Ab and PROTAC) following the rationale detailed in (github.com/chembl/tractability_pipeline_v2)^43^.

### High-throughput drug screens

For cell line drug screens, cells were dispensed into 384-well assay plates in 40 µl of growth medium using XRD384 (FluidX) dispensers. For organoid drug screens, organoids were cultured in 5% BME, then harvested, dissociated, counted and re-seeded in 5% BME suspension culture for 96 hours to allow organoid re-formation. Assay plates were then prepared with 10 µl of 50% BME:organoid medium using Multidrop Combi (Thermo Scientific) dispensers, and organoids from the re-formation cultures were harvested and transferred into the 384-well assay plates in 50 µl of their respective growth medium. For both model types, seeding densities had been optimised in advance to ensure cells remained in growth phase throughout the assay.

Following seeding, assay plates were incubated at 37 °C in a humidified atmosphere at 5% CO₂ for 24 hours, then dosed with the test compounds using an Echo555 (Labcyte) at a final DMSO concentration of typically 0.1% and treated for 72 hours. Viability was measured at the end of the assay by adding CellTiter-Glo 2.0 (Promega) to each well (10 µl for cell lines, 20 µl for organoids), followed by incubation at room temperature for 10 minutes. Luminescence was quantified using a Paradigm (Molecular Devices) plate reader. For each model, an untreated plate was read using CellTiter-Glo 2.0 at the time of dosing (i.e. 24 hours post-seeding) to provide the baseline signal used for growth estimates.

Assay plate quality control followed the approach previously described^132^, using the same coefficient of variation (CV) and Z-factor calculations. Plates were required to have a DMSO-treated negative control (NC-1) CV below 0.18 for cell lines and below 0.23 for organoids. Z-factors were calculated from NC-1 and each positive control (PC) (MG132, Staurosporine and medium-only blanks) and were required to exceed 0.3 for any control to which the model was sensitive (NC-1:PC ≥ 7). Vehicle effects were assessed as the ratio of untreated to DMSO-treated negative controls, required to fall between 0.8 and 1.2.

Compounds were screened as 7-point dose curves (seven concentrations per test compound, five concentrations per positive compound). Raw intensity data were analysed with the gdscIC50 R package. Data were normalised per plate to negative and positive controls, and dose-response curves were fitted using the non-linear mixed-effects model of Vis et al. (2016), as implemented in gdscIC50. All available replicates for a given model-compound pair contributed to a single fitted curve, from which IC50 and AUC were estimated. Curves with RMSE > 0.3 were excluded from downstream analysis. Drug targets and target pathways were further curated, and drugs tested in 12 or fewer models were removed.

Models are deemed sensitive to a given drug if their IC50 is less than or equal to the maximum concentration tested; otherwise, they are labelled resistant. A drug is classified as active across all models if the median IC50 is less than or equal to the maximum concentration tested; otherwise, it is classified as low-response. Drugs were compared across models using AUC. For each drug, the median AUC across models was calculated, and these values were used for comparisons across pathways.

### Biomarker analyses

We tested for associations between the response of models to CRISPR-Cas9 (n = 23) or drug screening (n = 36) and their underlying genomic features. Molecular features were derived from single-nucleotide variants (SNVs), short insertions/deletions (indels), copy number alterations (CNAs) and structural variants (SVs) in cancer driver genes or relevant PM genes. Driver events were grouped into three classes: mutation (Mut; nonsense, missense, stop-loss, frameshift and splice-site variants), amplification (Gain) and high-loss or homozygous deletion (Loss). For association testing, features were collapsed into two binary categories per gene, mutated/lost (Mut/Loss) and Gain, reflecting the expected direction of functional consequence. Genes that were recurrently co-altered across the panel, or paralogs (e.g., LATS1/LATS2) were merged into composite features, with a model annotated as altered if it carried at least one qualifying alteration in any of the constituent genes. A gene-level feature was retained for testing only if it was altered in at least three models in the CRISPR screen cohort, or four models in the drug screen cohort.

To select CRISPR-Cas9 genes for further analysis, we first computed the median expression of each gene across all PM models, and restricted testing to genes with a median log₁₀(TPM + 1) ≥ 1. Genes were further required to show a dependency in at least three models and no dependency in at least three models (n=2035). For the drug analysis, we used the 36 models with available drug screening data and restricted testing to the 82 compounds with at least three sensitive models.

To test associations between genomic features and the previously defined phenotypic responses, we used the analytical framework implemented in GDSCTools v0.17^133^ (github.com/CancerRxGene/gdsctools), previously applied to large-scale pharmacogenomic and fitness datasets^9,10^. For each gene feature-response pair, we fitted an ordinary least-squares model of the form: *Y* ∼ *C*(*Model*_*Type*) + *Mutation*_*Feature*, where the response Y is the scaled depletion log fold change for CRISPR-Cas9 associations, or the log₁₀(IC₅₀) for drug associations, C(Model_Type) is a categorical covariate encoding model type (organoid or cell line), and Mutation_Feature is the gene-level feature under analysis. P values from tested associations were corrected using the qvalue method implemented in GDSCtools, applied at local level, and associations with p-value < 0.05 and FDR < 25% were considered significant and were selected for manual review. Associations were classified as sensitising or conferring resistance according to the signed effect size.

## Data availability

The raw WGS and RNA-seq data generated in this study will be deposited in the European Genome-Phenome Archive (EGA). The raw WGS and RNA-seq data from primary cell lines used in this study have been deposited in the European Genome-Phenome Archive (EGA): WGS, EGAD00001009641; RNA-seq, EGAD00001009642. Processed data files and links to raw data files are available at Cell Model Passports (CMP) (cellmodelpassports.sanger.ac.uk/) or available as supplementary files. Additional supplementary files and code will be available on Zenodo. The PM organoid models will be made available for research purposes.

## Acknowledgments

This work was funded by Mick Knighton Mesothelioma Research Fund-British Lung Foundation Mesothelioma Programme Grant (ABR 12-1) and Project Grant MKMRFPG18-3 and the June Hancock Mesotheloma Research Fund. This work was funded, in part, by Wellcome Trust Grant 206194 and 220540/Z/20/A. RCR was funded by the NIHR Cambridge Biomedical Research Centre (NIHR203312) and Cancer Research UK Cambridge Centre (CTRQQR-2021\100012). J.A.V. is supported by Asthma & Lung UK grant MPG24\11. We thank Professor Marion MacFarlane and Dr Zsuzanna Tabi for their valuable contributions to the derivation of the primary PM cell lines used in this study^28,134^. We would like to acknowledge colleagues at the Wellcome Sanger Institute: the Translational Cancer Genomics group for generating the drug screening data and analyses; Cellular Operations for deriving the organoid models and performing CRISPR screening on them; and the Somatic Genomics Programme Laboratory Team for downstream pellet processing.

For the purpose of Open Access, the author has applied a CC BY public copyright licence to any Author Accepted Manuscript version arising from this submission.

## Extended Figure Legends

**Extended Figure E1. Multimodal profiling of low passage PM primary cell lines.** (A) Schematics of the expansion workflow and the integrated dataset generated (clinical annotation, WGS, RNA-seq, CRISPR–Cas9 dependency screening and high-throughput drug screening). (B) Cohort characteristics and sample attributes, including sex, age, histological subtype. (C) Data completeness across the 21 PM primary cell lines for molecular profiling and functional screens.

**Extended Figure E2. Comparison of genomic features between PM organoids and matched tumours.** (A) Per-model tumour–organoid concordance for purity and fraction of shared mutations, together with a landscape of recurrent driver alterations present in the matched tumour (T) and organoid (O). Driver alterations are shown with event type. (B) Matched tumour–organoid fraction of shared mutations across the whole genome (n=26) (left). Correlation of fraction of shared mutations with tissue purity (right). (C) Clonal fraction of unique tumours (n=23) and organoids (n=24) (left), and correlation of clonal fraction in matched tumours and organoids (right).

**Extended Figure E3. Mutational signatures analysis of hypermutant cases and PM cohorts.** Comparison of SBS96 (A) and ID83 (B) profiles across unique PM organoids (n=23), PM matched tissues (n=22), and TCGA samples (n=69) (left subplots), or across hypermutant PM organoid, hypermutant matched tissue, and hypermutant case in TCGA cohort (right subplots). (C) Exome-based SBS96 and ID83 (D) mutational profiles of hypermutant organoid (WTSI-MESO_098), its matched tissue, and TCGA hypermutant case (left). Exome-based mutational profiles of aggregated PM organoid cohort, their matched tissues, and TCGA cohort, all excluding hypermutant cases (right).

Extended Figure E4. CRISPR fitness genes per model. (A) Bar plot (top) showing total number of fitness genes per PM organoid model. Boxplot (right) shows the distribution of fitness genes across PM organoids (n=14). Fitness effect (LFC scaled) across all tested genes splitted by essential (coloured) or non-essential (grey) per model (bottom). Aggregated fitness effect across PM organoids for identified essential and non-essential genes (right). (B) Bar plot (top) showing total number of fitness genes per primary PM cell line model. Boxplot (right) shows the distribution of fitness genes across PM cell lines (n=9). Fitness effect (LFC scaled) across all tested genes splitted by essential (coloured) or non-essential (grey) per model (bottom). Aggregated fitness effect across PM cell lines for identified essential and non-essential genes (right). (C) Number of fitness genes identified as essential in at least N number of models or more (x-axis). The red dashed line depicts the threshold for PM core fitness genes.

## Supplementary Figure Legends

**Supplementary Figure S1. PM organoid derivation protocols.** Schematic representation of PM organoid derivation protocols from primary tissue (A) or pleural fluid (PM) until seeding. Step-by-step detailed information in Methods.

**Supplementary Figure S2. Histopathological analyses.** Representative images of organoids and parental tumours (tissue) for haematoxylin and eosin (H&E), Mesothelin, Podoplanin, Calretinin, Cytokeratin 6/5, and Wilms tumour (WT1). Where parental tumour tissue was available, histological images are displayed adjacent to the corresponding organoid. Different models derived from matched pleural fluid and primary tumour from the same donor are shown as pairs. Scale bars are 50 μm.

**Supplementary Figure S3. Genome-wide tumour-organoid SCNA comparisons.** Genome-wide somatic copy-number alteration (SCNA) profiles for organoids (orange) and matched tumours (purple). Each dot corresponds to a 100kbp segment. Pearson correlation R and p-value are annotated in the title of each subplot.

**Supplementary Figure S4. Comparison of banked organoids with matched early-passage versions.** (A) Per-model comparison of banked organoids (organoid, orange) and their early versions (early organoid, yellow) for purity, ploidy, mutational burden and fraction of shared mutations, together with a landscape of recurrent driver alterations. Driver alterations are shown with event type. (B) Matched fraction of shared mutations across the whole genome (n=8) (left). Matched SCNA concordance (Pearson R), showing high genome-wide similarity (n=6) (right). (C) Genome-wide somatic copy-number alteration (SCNA) profiles for organoids (orange) and their early passage version (yellow). Each dot corresponds to a 100kbp segment. Pearson correlation R and p-value are annotated in the title of each subplot. (D) Per-sample relative mutation type comparing matched early-version (1) and banked organoid (2) pairs, stratified by mutation class (SNVs vs indels). Bar plots additionally show relative contributions of single-base substitution (SBS) and indel (ID) mutational signature activities, and their grouped profiles (right). (E) Cosine similarities between pairs based on SBS96 and ID83 mutational profiles, demonstrating high concordance across all sample pairs. (F) Aggregate SBS and ID mutational profiles used for signature extraction, cosine similarity detailed in the titles.

**Supplementary Figure S5. Primary PM cell lines whole-genome sequencing and their genomic landscape.** (A) Per-model clinical annotation, mutational burden, ploidy, and purity, together with a landscape of recurrent driver alterations. Driver alterations are shown with event type, alongside summary bar plots of alteration types and number of altered cell lines (right). (B) Genome-wide somatic copy-number alteration (SCNA) profiles for primary PM cell lines, with key loci indicated (e.g., for example SETD2, BAP1 and others), illustrating preservation of hallmark pleural mesothelioma copy-number patterns. (C) Comparison between mutational burden of primary PM cell lines (n=21) and established PM cell lines (n=25). (D) Clonal fraction of primary PM cell lines (n=21).

**Supplementary Figure S6. Genome-wide primary PM cell lines SCNA comparisons.** Genome-wide somatic copy-number alteration (SCNA) profiles for primary PM cell lines (green). Each dot corresponds to a 100kbp segment.

**Supplementary Figure S7. Mutational profile pairs for SNV.** SBS96 mutational profiles of organoid (upper) and matched tumour (lower) pairs, cosine similarities detailed in title.

**Supplementary Figure S8. Mutational profile pairs for indels.** ID83 mutational profiles of organoid (upper) and matched tumour (lower) pairs, cosine similarities detailed in title.

**Supplementary Figure S9. Transcriptomic relationships among mesothelioma models.** (A) Sample-sample Pearson correlation R heatmap of variance-stabilised, normalised expression counts across the 500 most variable protein-coding genes. Samples are ordered by hierarchical clustering and annotated by model type and histological subtype. Colour intensity (purple scale) indicates Pearson correlation coefficients R. (B) Principal component analysis (PCA) based on the 500 most variable genes. Samples are coloured and shaped according to model type (organoid or primary cell line) and histological subtype (epithelioid or non-epithelioid, including biphasic and sarcomatoid), as indicated in the legend.

**Supplementary Figure S10. CRISPR-Cas9 screening quality control.** PM organoids (n=15, before filtering) (A) and PM primary cell lines (n=9) (B). Distributions of mean Euclidean distances across replicate comparisons for each model. Note the organoid WTSI-MESO_009 failed QC and was discarded (left). AUROC and AUPR curves for each model (grey) and the mean per model type (colour) (centre). Fitness effect (scaled LFC) for reference essential and reference non-essential genes across all models; the distribution of reference is centred at −1, and non-essential at 0 (right).

**Supplementary Figure S11. Summary of tractability data for fitness genes.** (A) Fitness genes depleted in N models (x-axis) classified according to the tractability classification for small molecules (SM), antibodies (Ab), and PROTACs. Colour scale indicates the number of genes per category as shown in each square. Further classification in tractability buckets for SM (B), Ab (C), and PROTAC (D), following the same format detailed in A.

**Supplementary Figure S12. Fitness genes tractability prioritisation for antibodies and PROTACs.** Identified priority targets were classified according to Antibodies (A) and PROTACs (B) tractability evidence and assigned to tractability buckets and categories as detailed in each plot. Colour scale indicates the number of models in which the gene is essential; text colour indicates the broad pathway or functional group of the gene, y-axis indicates the median of the negative fitness effect across models where the gene was considered essential; a higher y-axis indicates a stronger fitness effect.

**Supplementary Figure S13. Drug responses across PM models.** (A) Heatmap showing the response of each PM model to each drug in the drug screen panel. Sensitive models to a drug are shown in blue, resistant in red. Models are ordered by hierarchical clustering, applied separately to organoids and cell lines. (B) Heatmap showing the resulting AUC for each model-drug assay. Models are sorted as in A.

## Supplementary Table Legends

**Supplementary Table 1.** Clinicopathological features of donors in the PM organoid cohort and data availability.

**Supplementary Table 2.** PM organoid derivation attempts and success status.

**Supplementary Table 3.** Clinical annotation of low passage PM primary cell lines and data availability.

**Supplementary Table 4.** Summary genomic characterisation of PM models and tissues.

**Supplementary Table 5.** Summary of driver alterations across PM organoids.

**Supplementary Table 6.** Summary of driver alterations across primary PM cell lines.

**Supplementary Table 7.** Activity of SNV (SBS96) mutational signatures across PM organoid models and matched tissues.

**Supplementary Table 8.** Activity of indels (ID83) mutational signatures across PM organoid models and matched tissues.

**Supplementary Table 9.** Differential expression analysis between PM organoids and PM primary cell lines.

**Supplementary Table 10.** Functional enrichment analysis of differentially expressed genes between PM organoids and PM primary cell lines.

**Supplementary Table 11.** Fitness genes and fitness effect per PM model.

**Supplementary Table 12.** PM core fitness genes.

**Supplementary Table 13.** Functional enrichment analysis of PM core fitness genes.

**Supplementary Table 14.** Tractability assessment of fitness genes across PM models.

**Supplementary Table 15.** Genomic features and CRISPR-Cas9 associations.

**Supplementary Table 16.** Drug responses across PM models.

**Supplementary Table 17.** Genomic features and drug response associations.

