## Supplementary Figures for "Multiomic analysis of a pleural mesothelioma organoid biobank defines gene dependencies and therapeutic vulnerabilities"

### Supplementary Figure S1

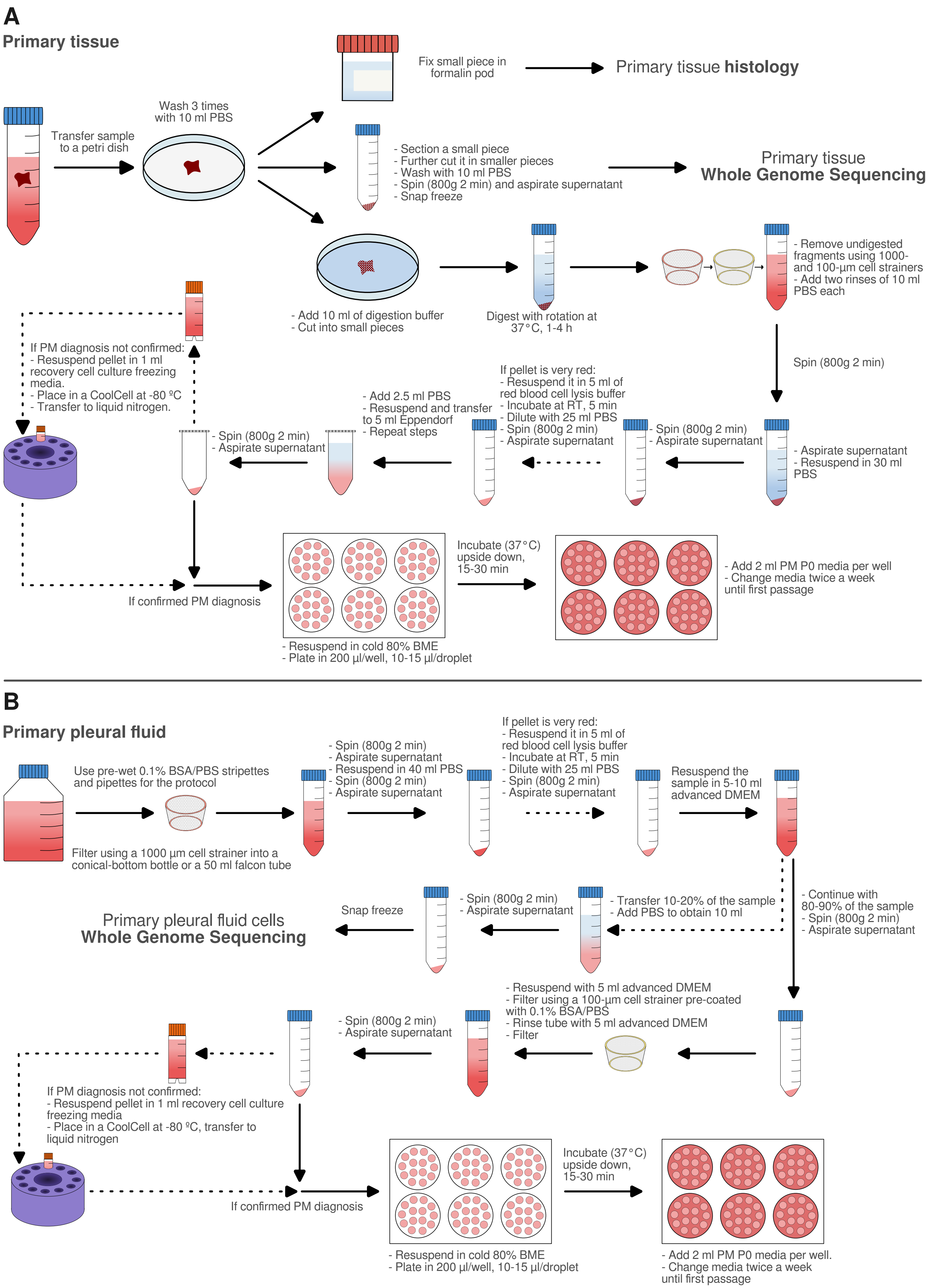

### Supplementary Figure S2_a

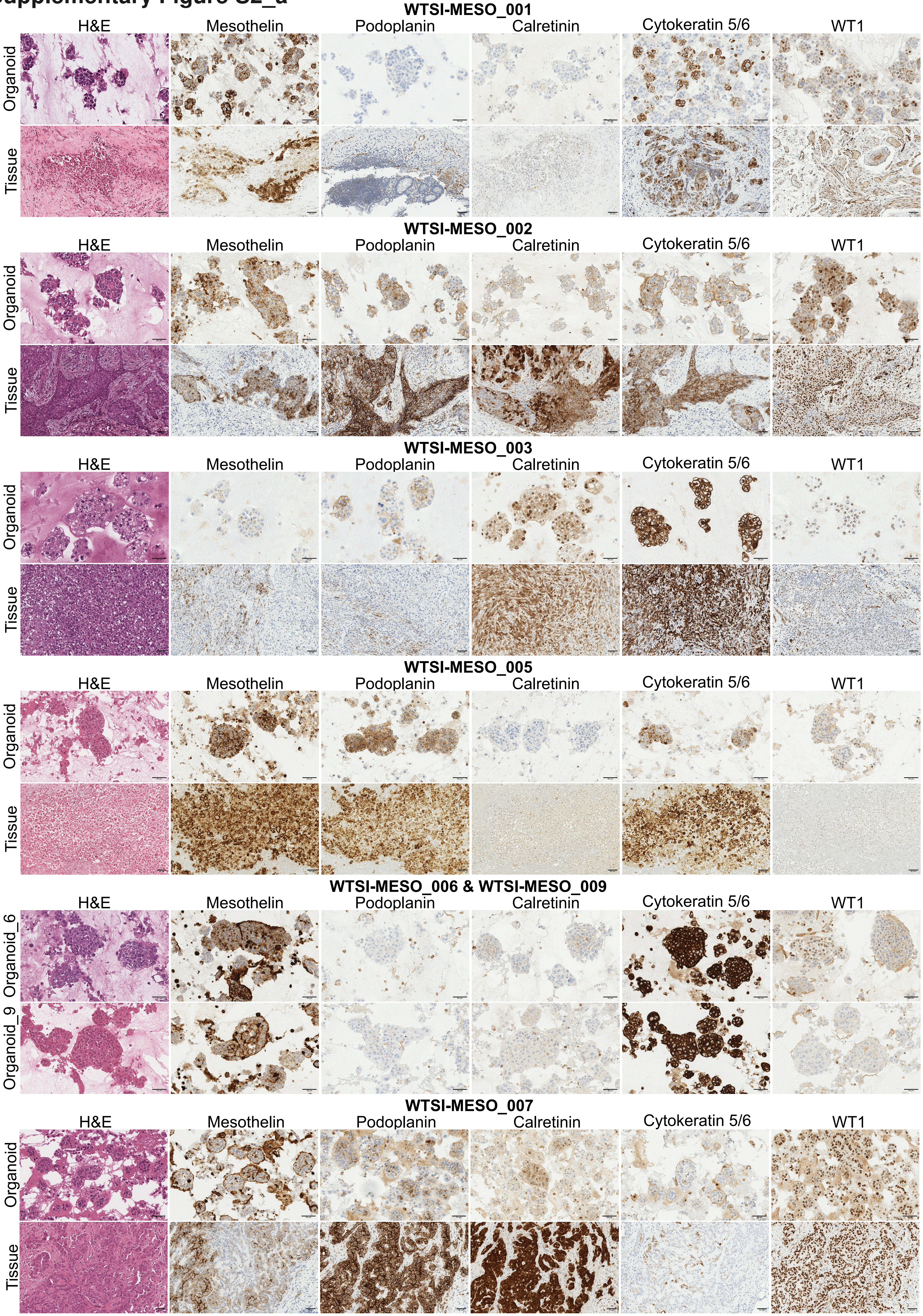

### Supplementary Figure S2_b

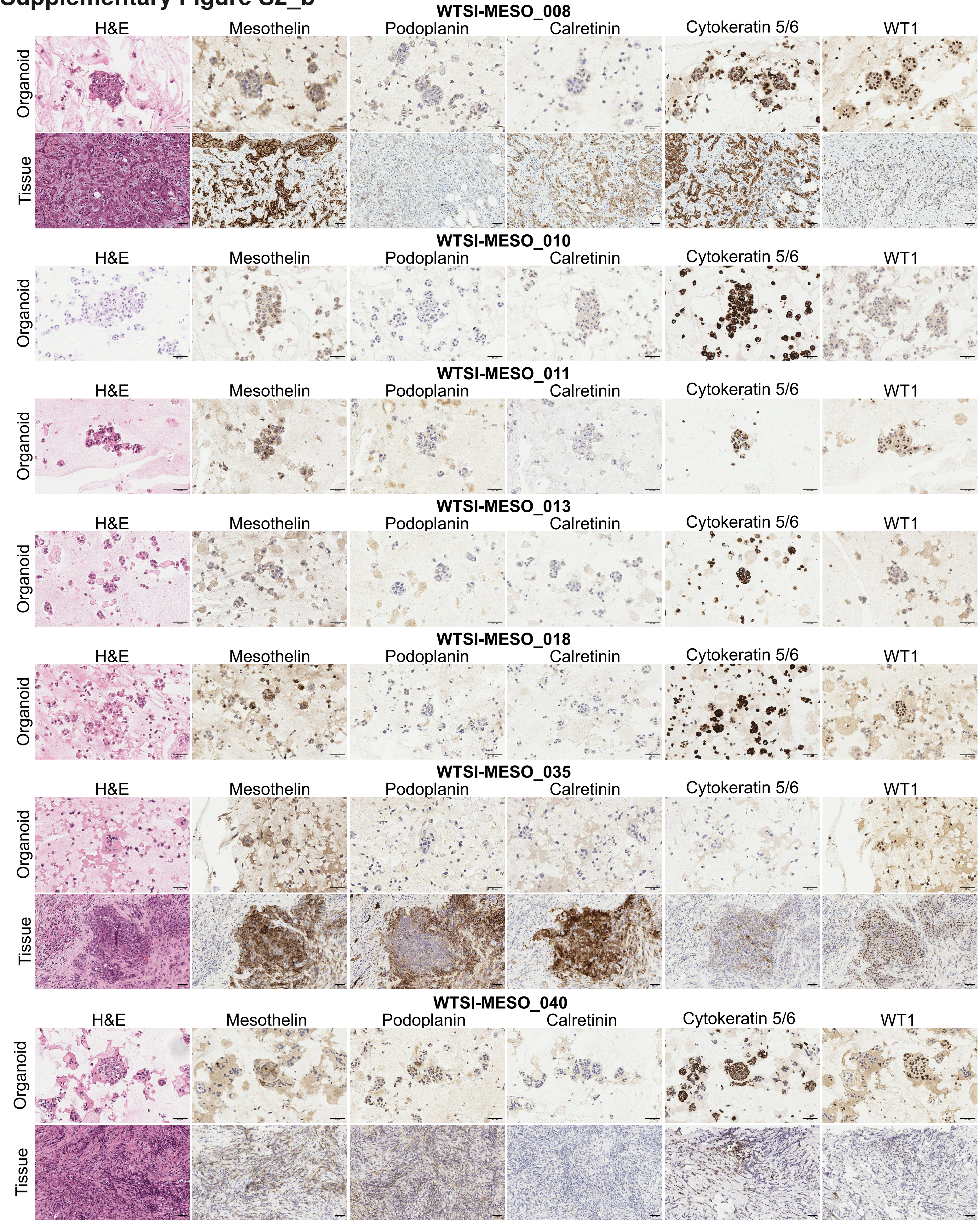

### Supplementary Figure S2_c

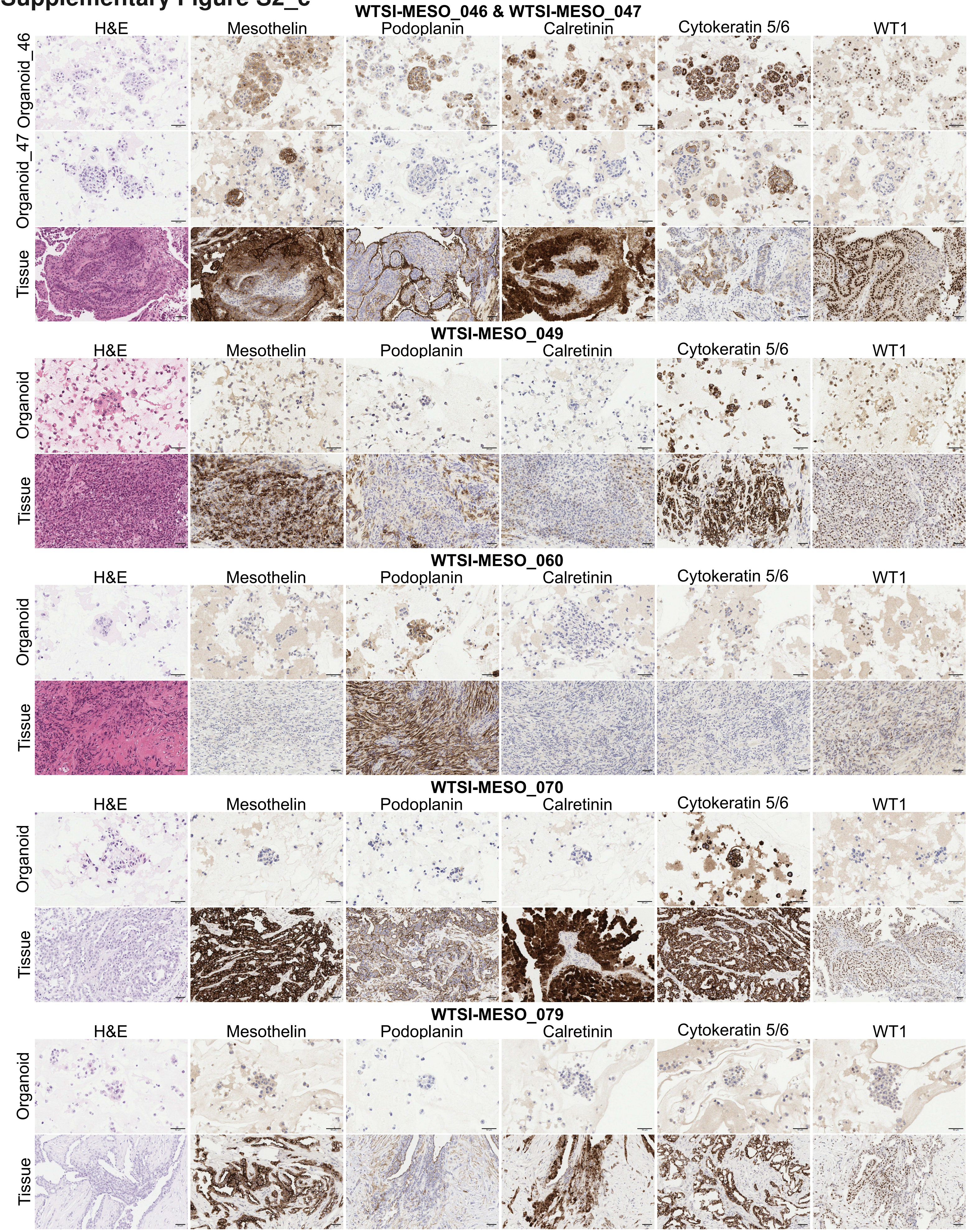

### Supplementary Figure S2_d

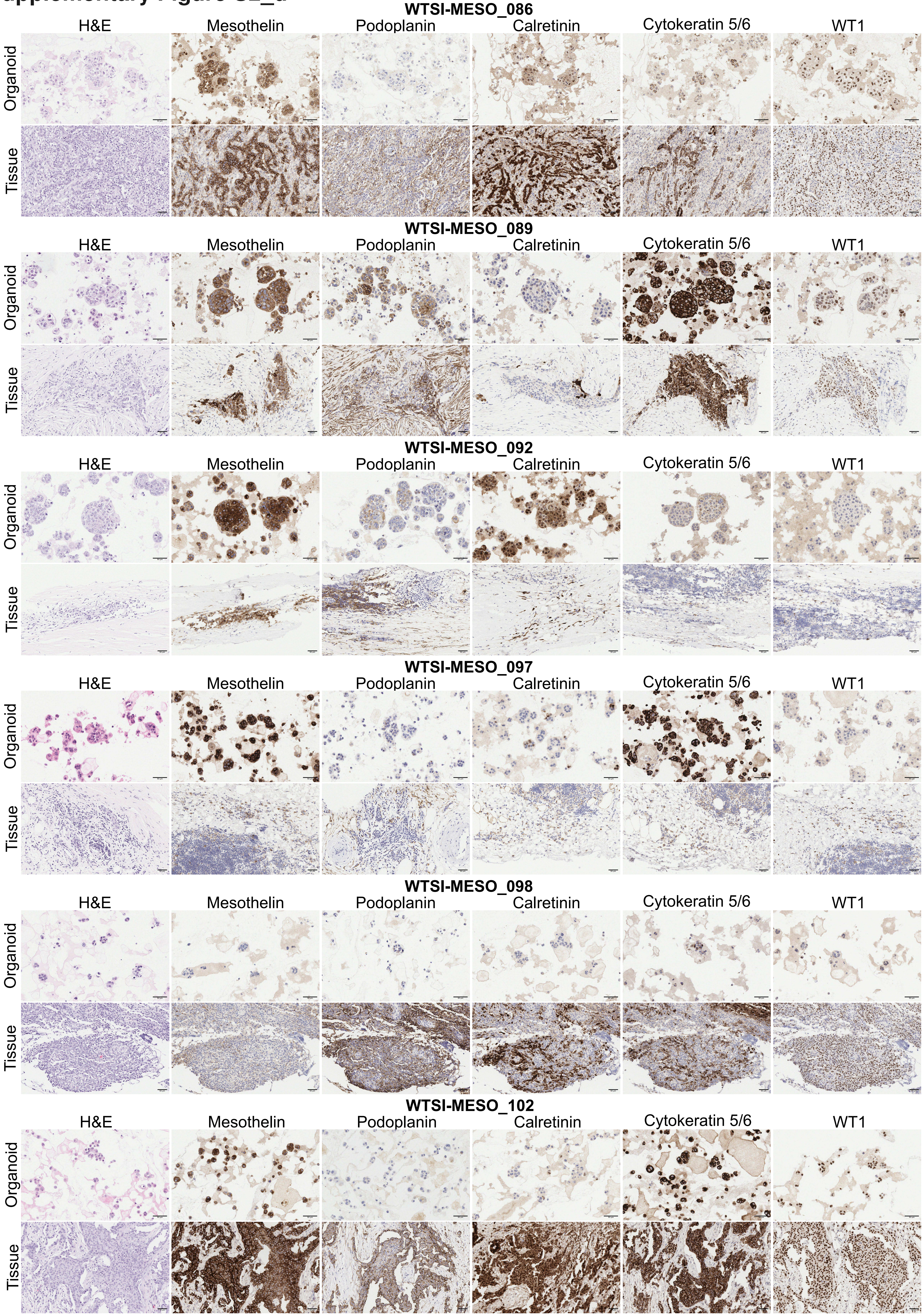

### Supplementary Figure S3

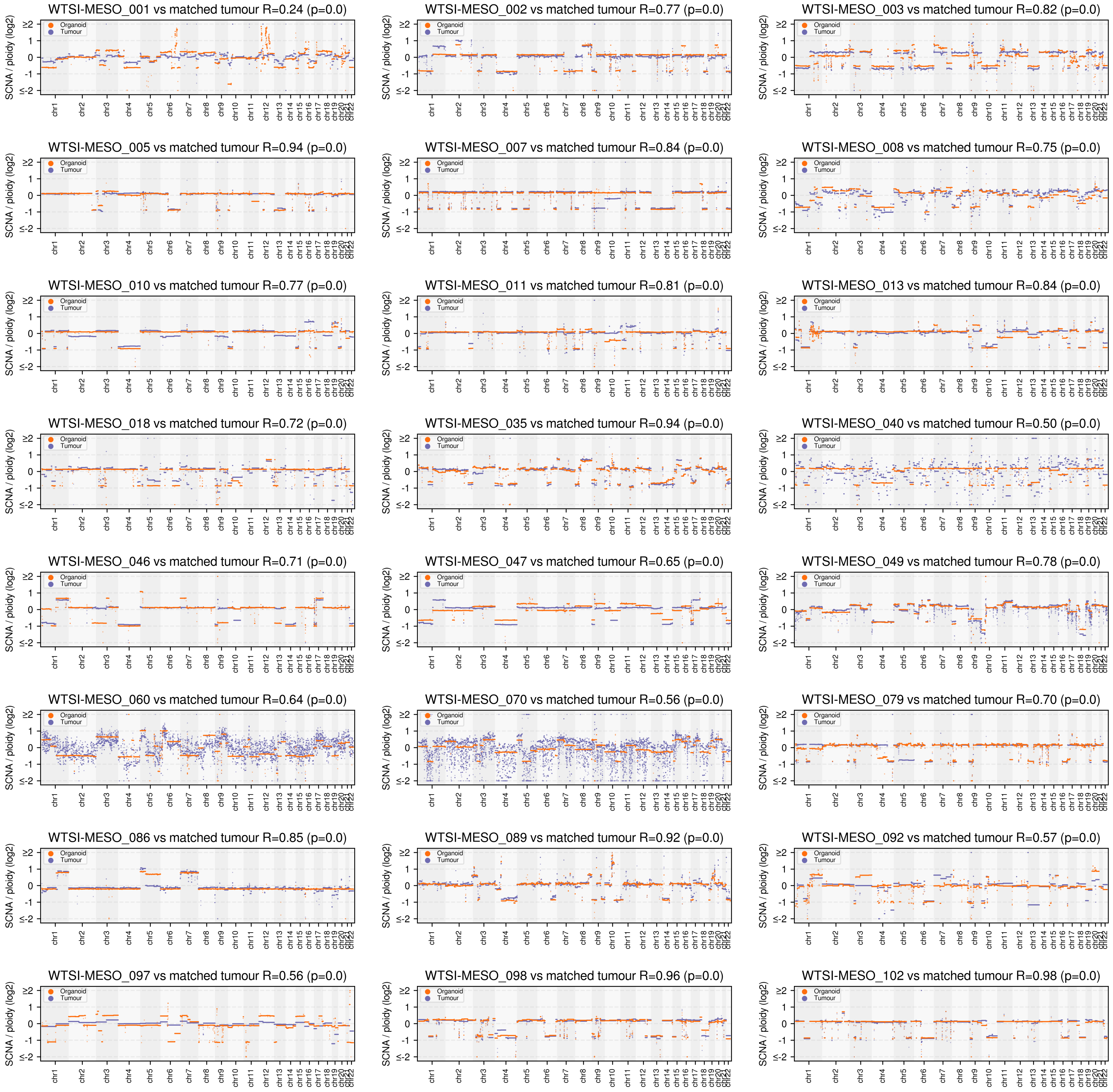

### Supplementary Figure S4

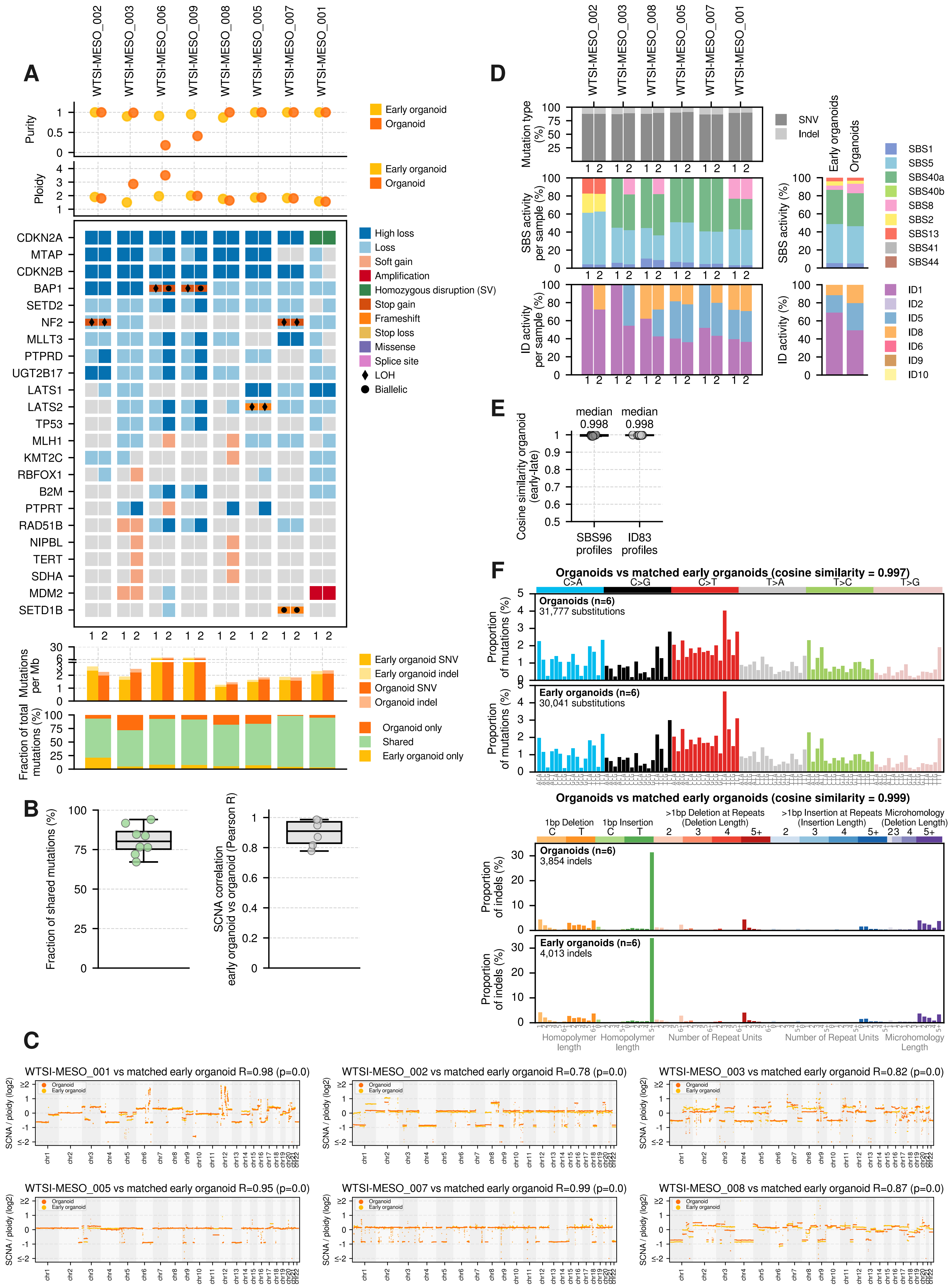

### Supplementary Figure S5

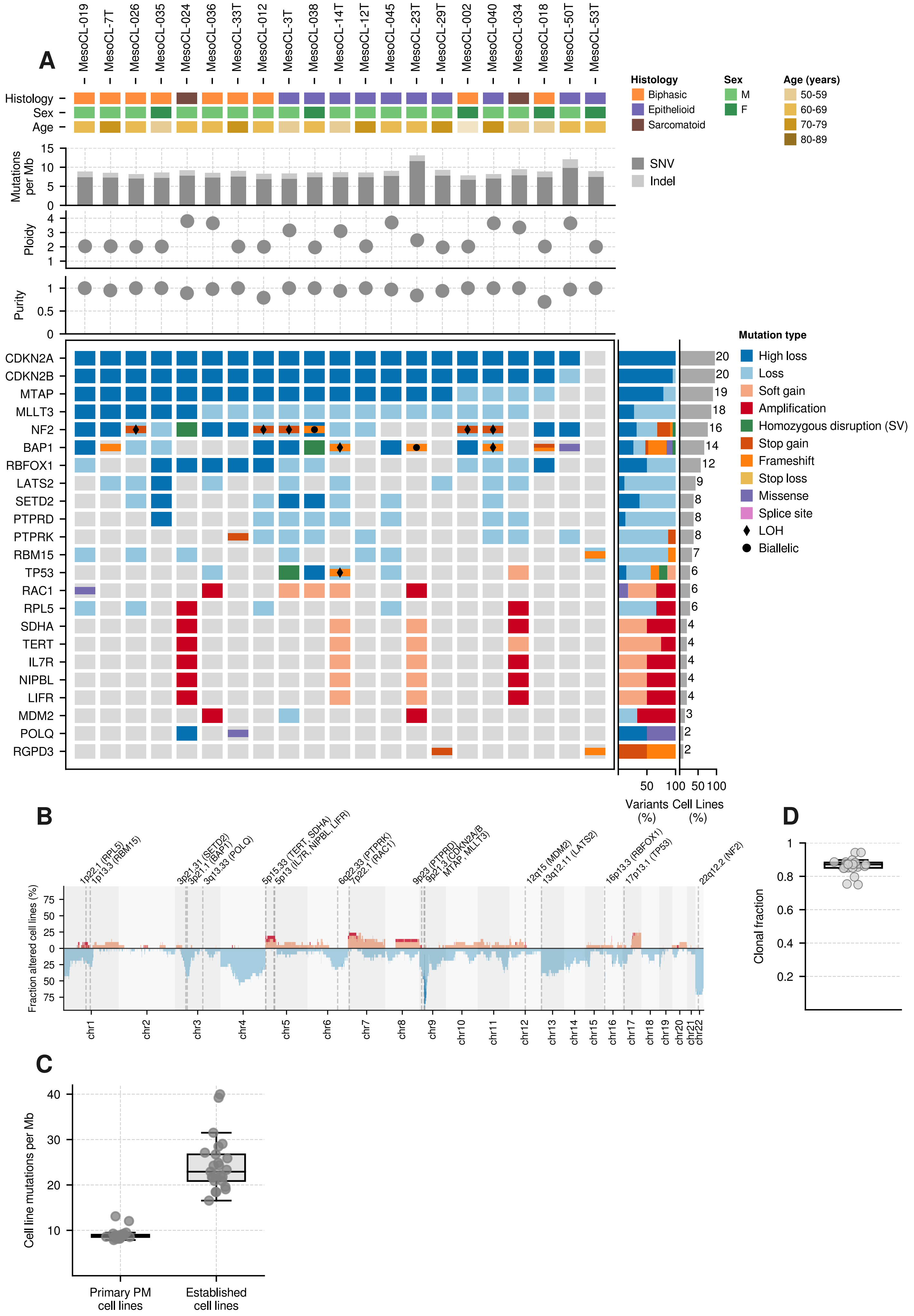

### Supplementary Figure S6

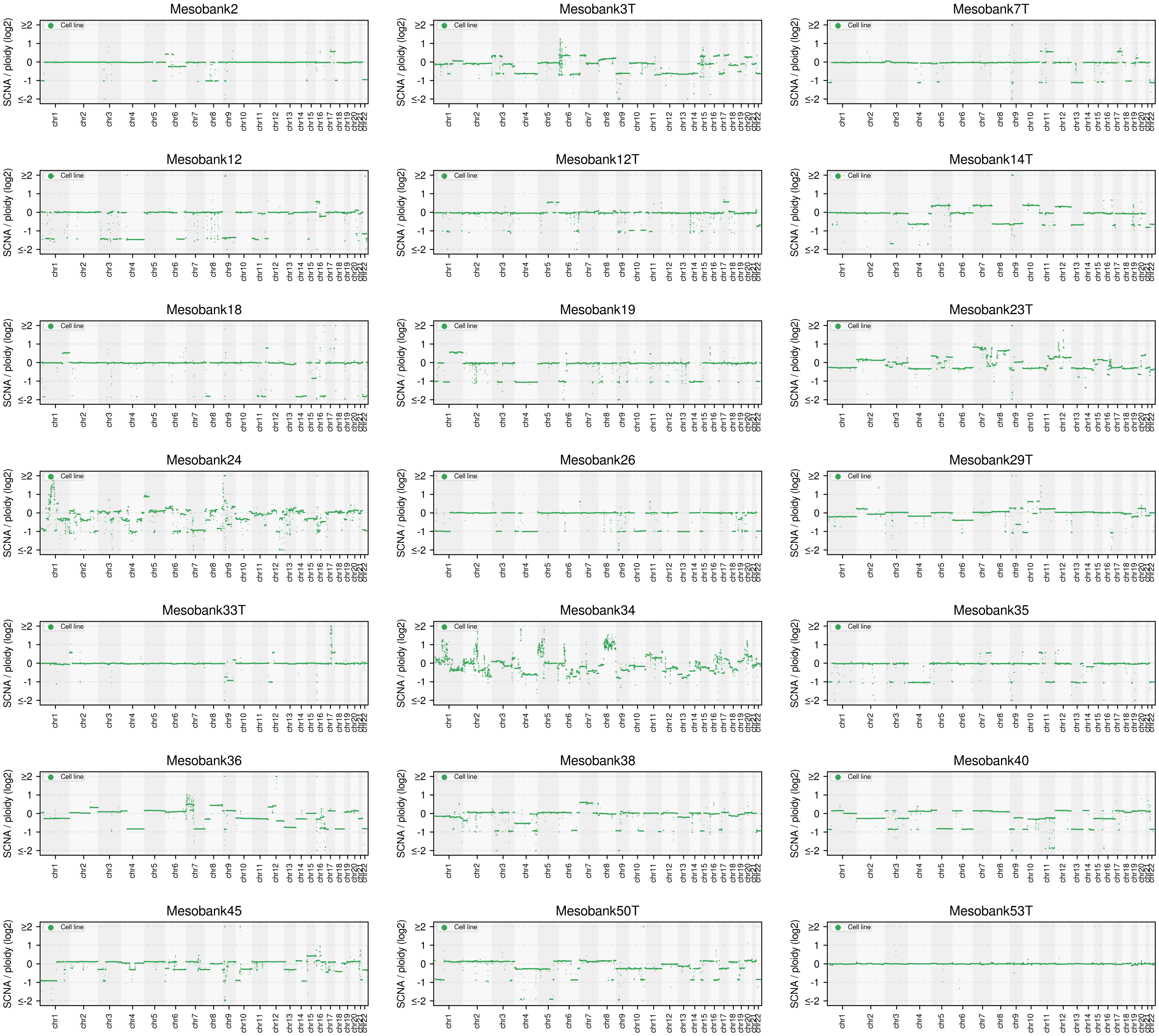

### Supplementary Figure S7

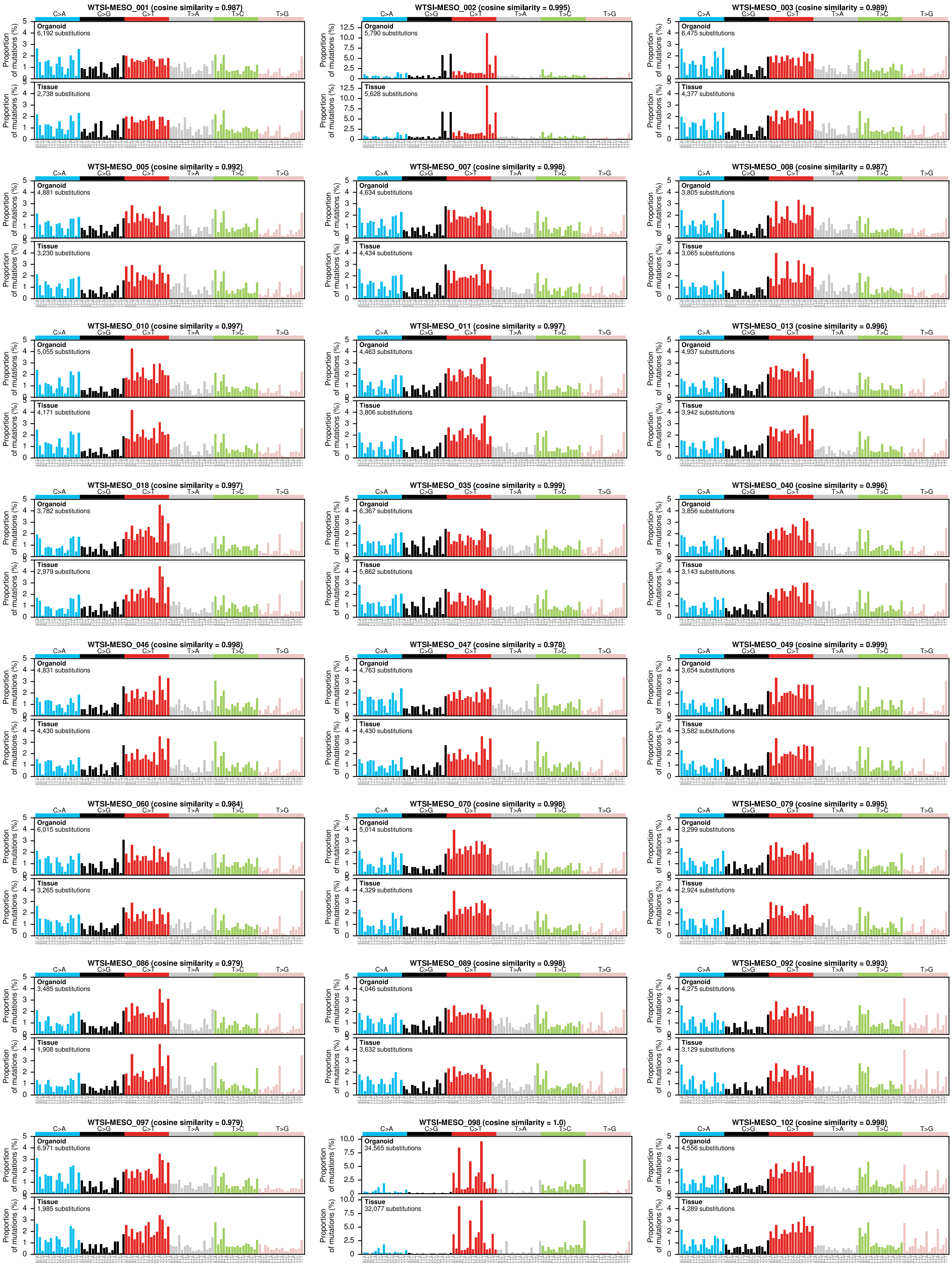

### Supplementary Figure S8

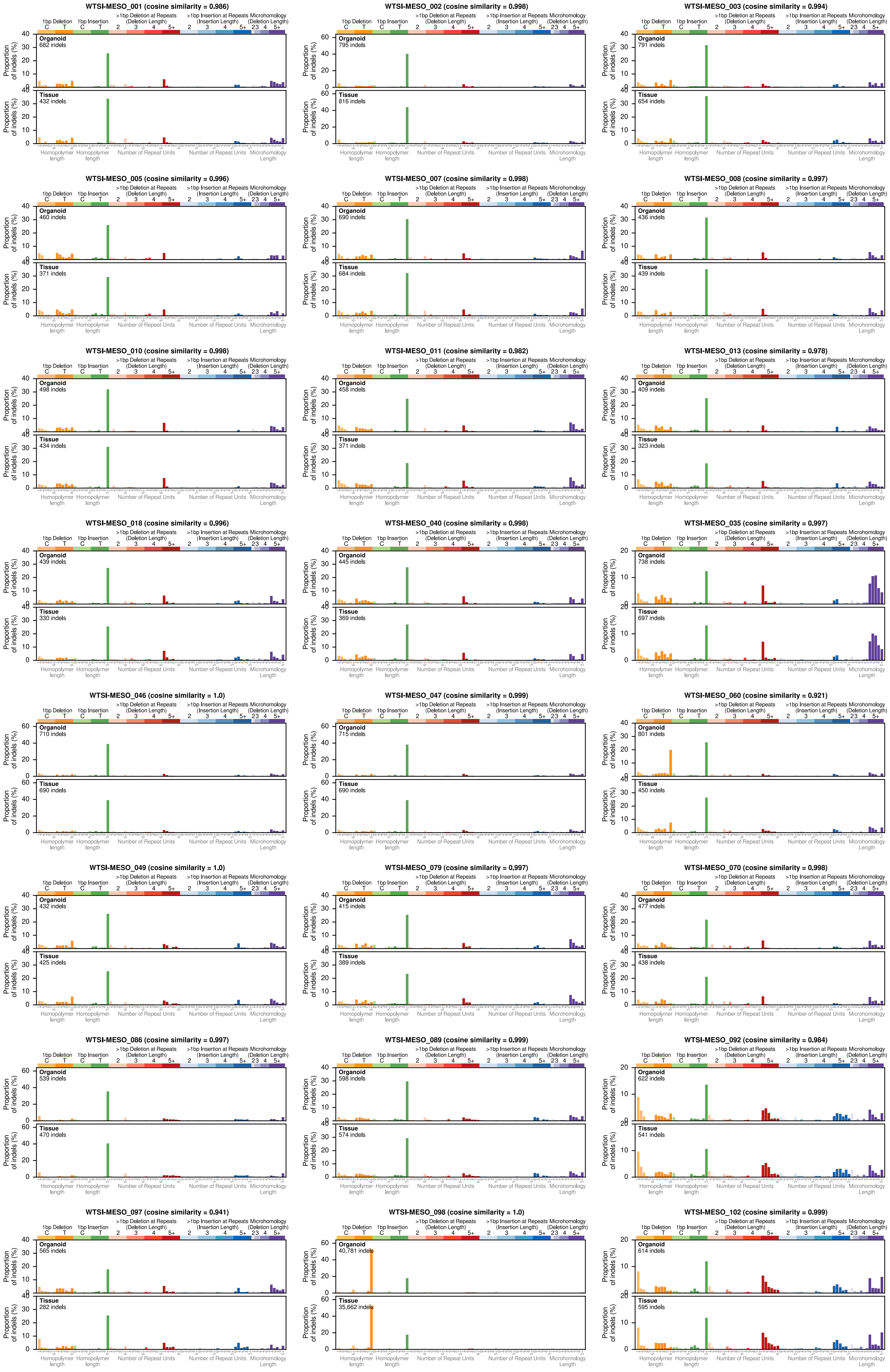

### Supplementary Figure S9

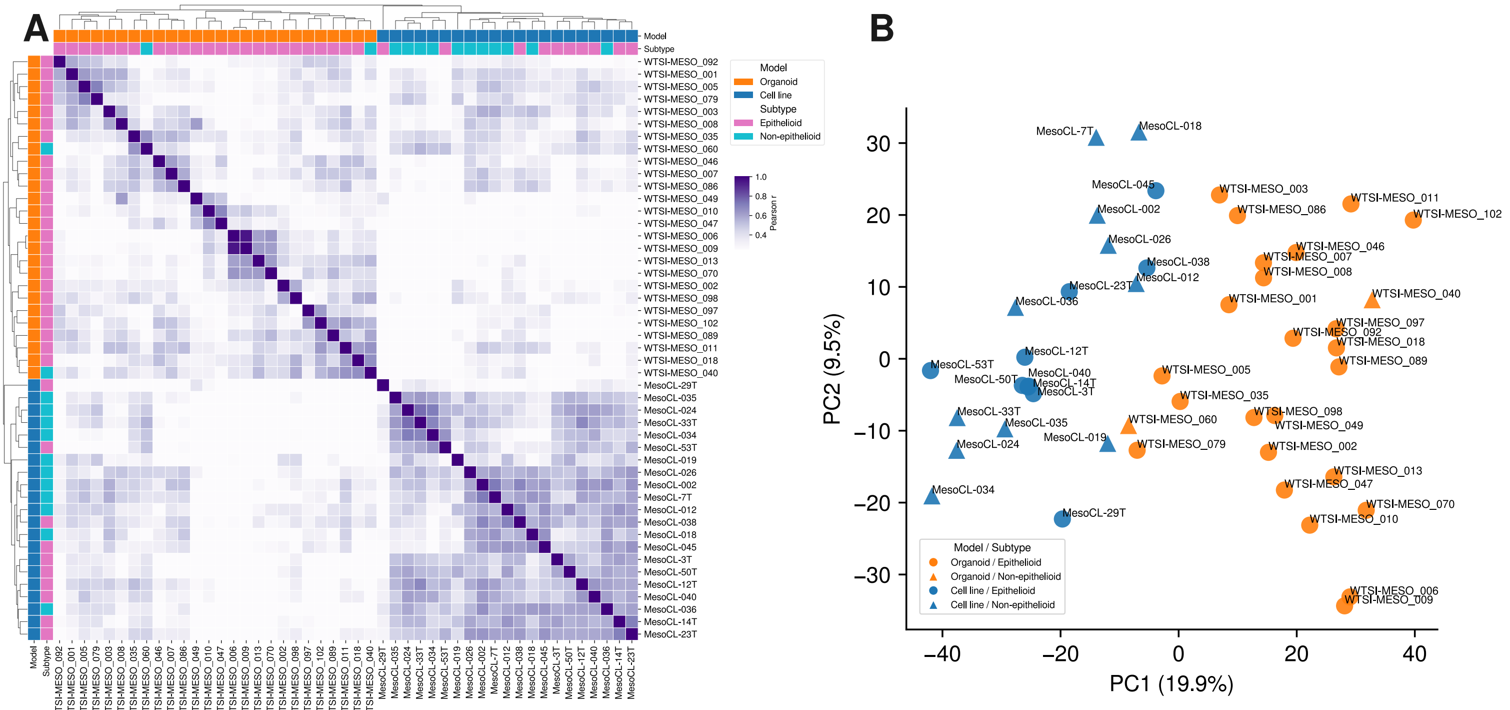

### Supplementary Figure S10

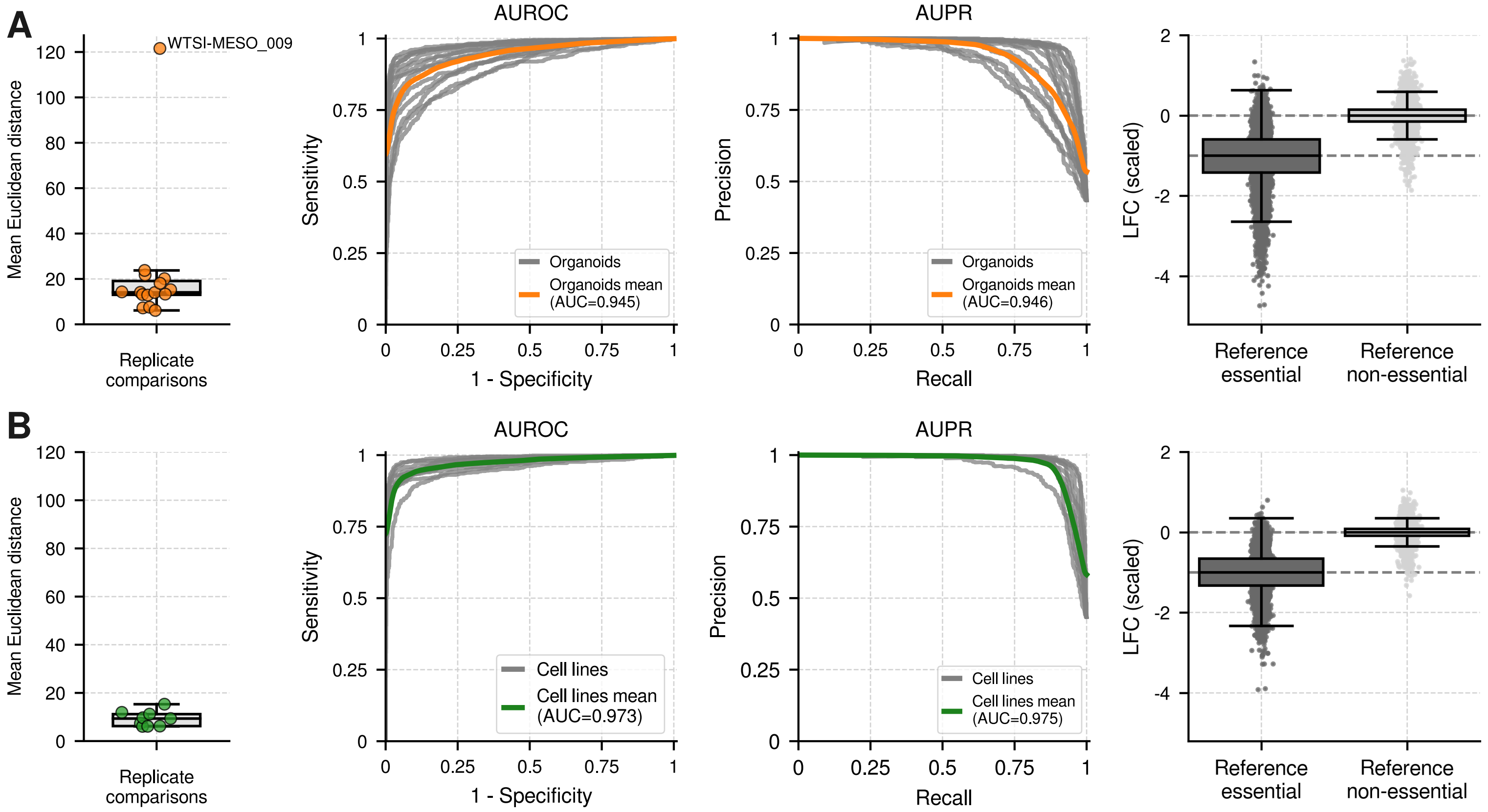

### Supplementary Figure S11

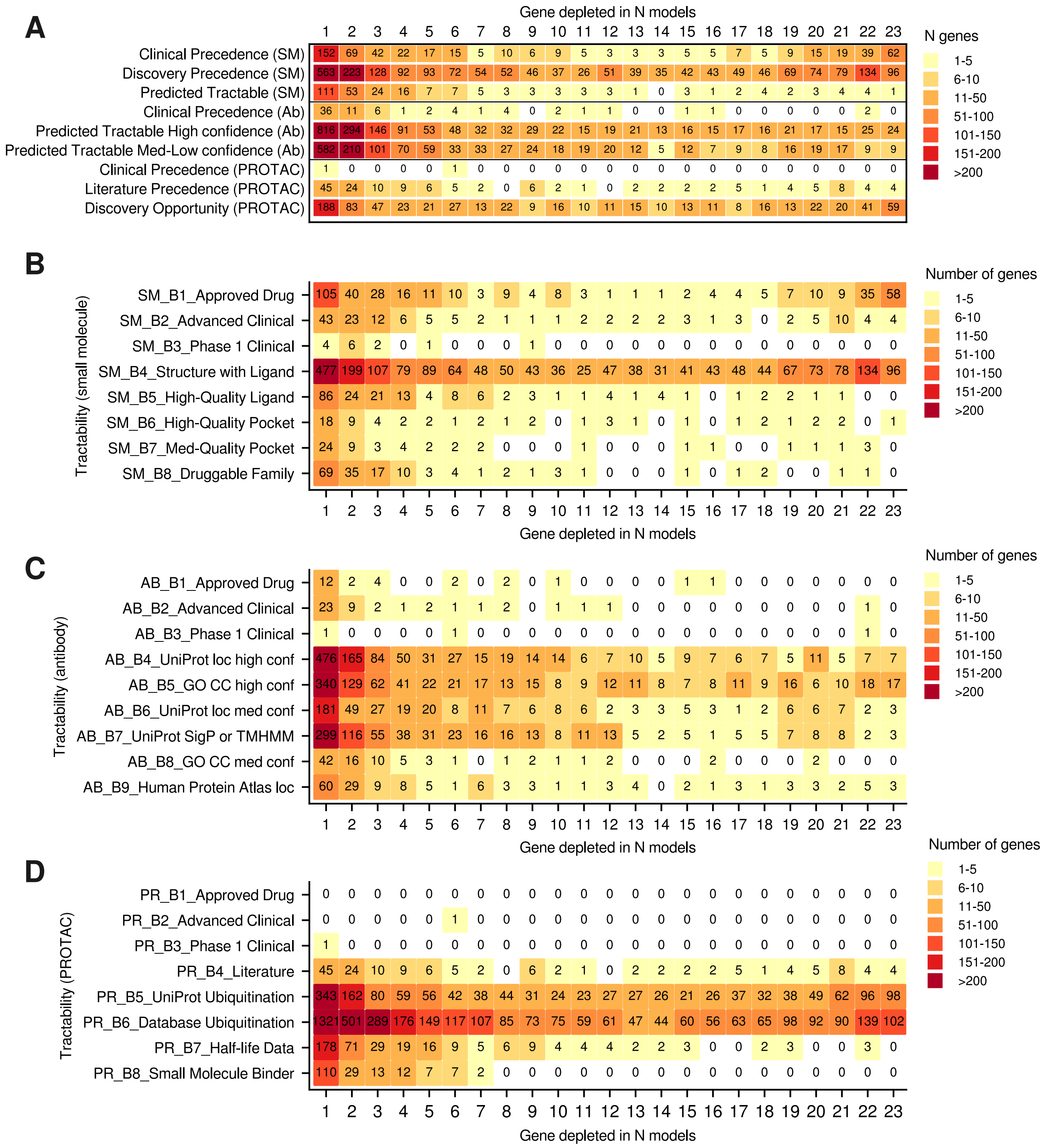

### Supplementary Figure S12

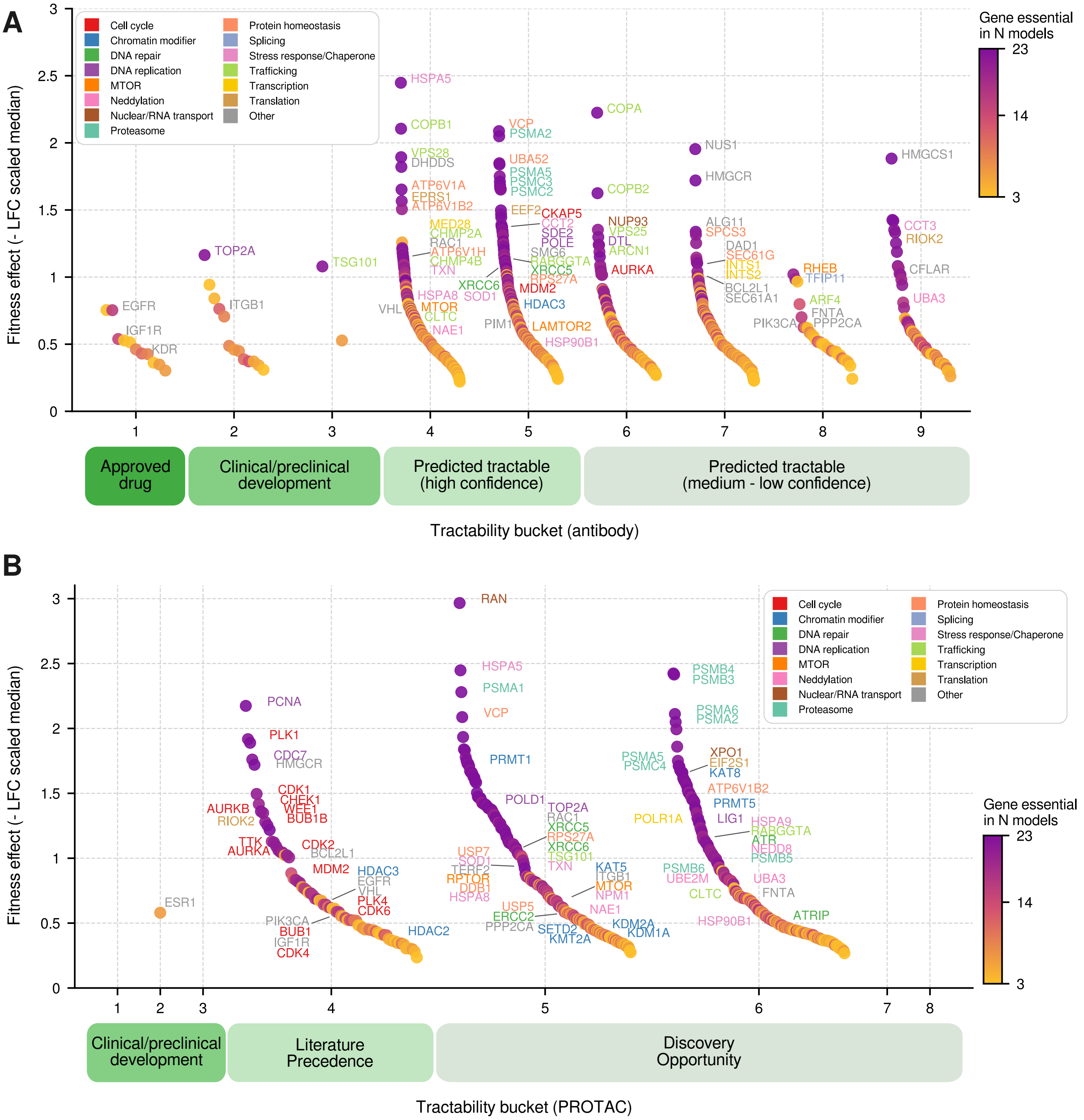

### Supplementary Figure S13

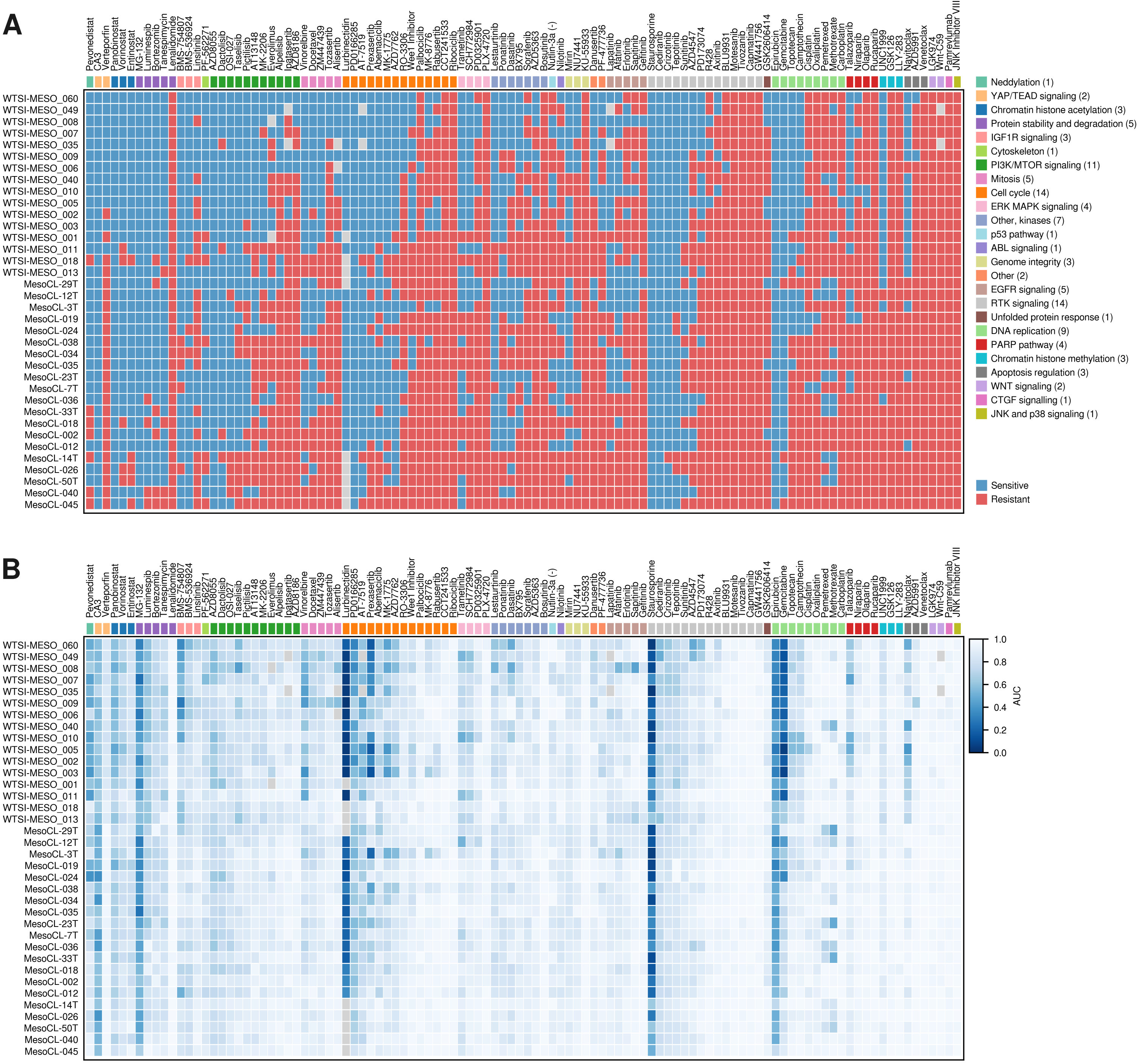
