## Supplementary figures and images for "Multiomic analysis of a pleural mesothelioma organoid biobank defines gene dependencies and therapeutic vulnerabilities"

### Extended Figures

#### Extended Figure E1

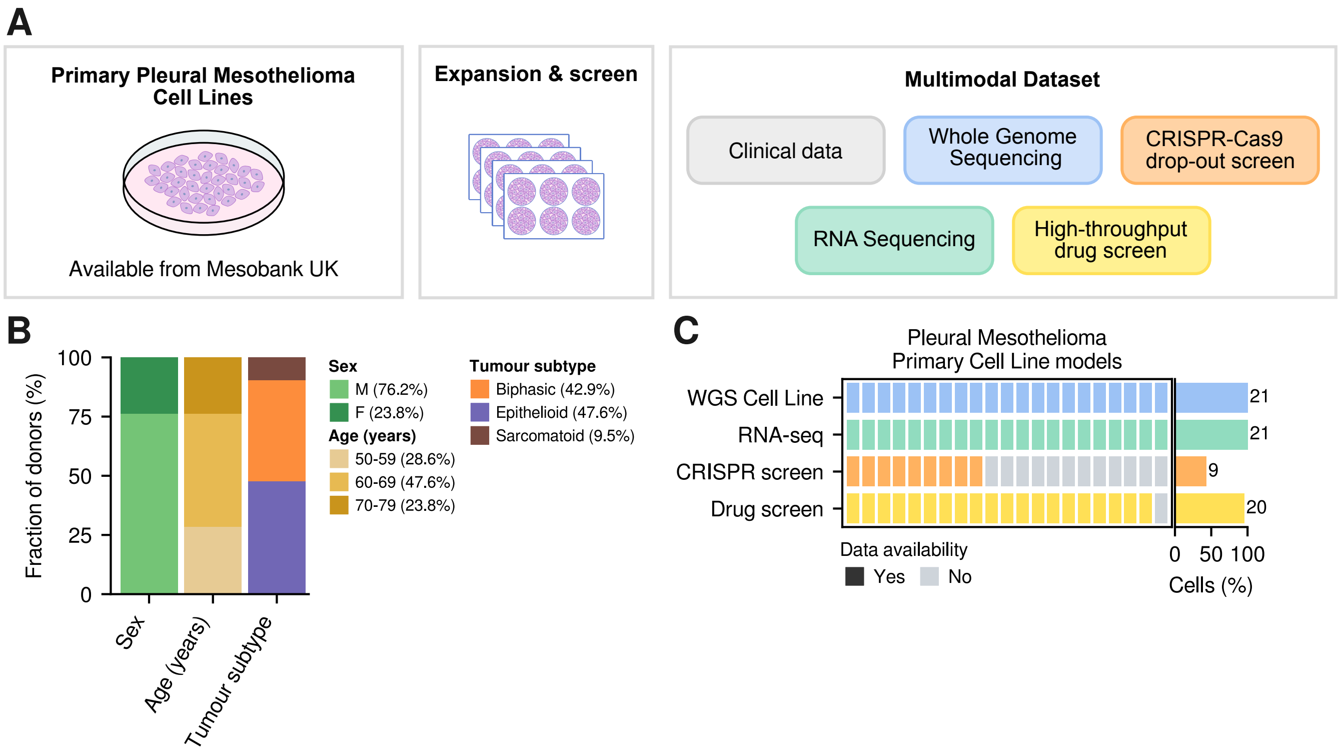

#### Extended Figure E2

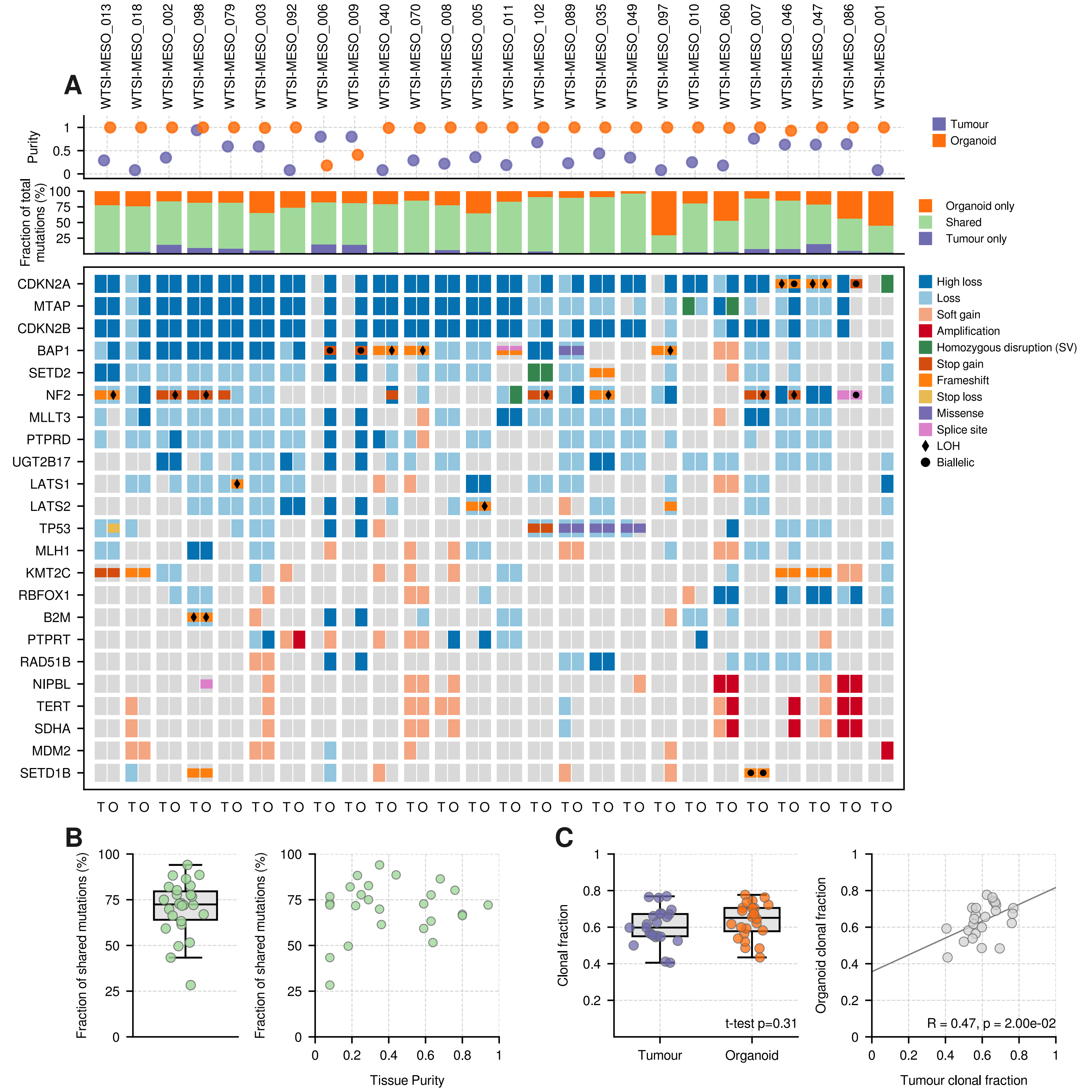

#### Extended Figure E3

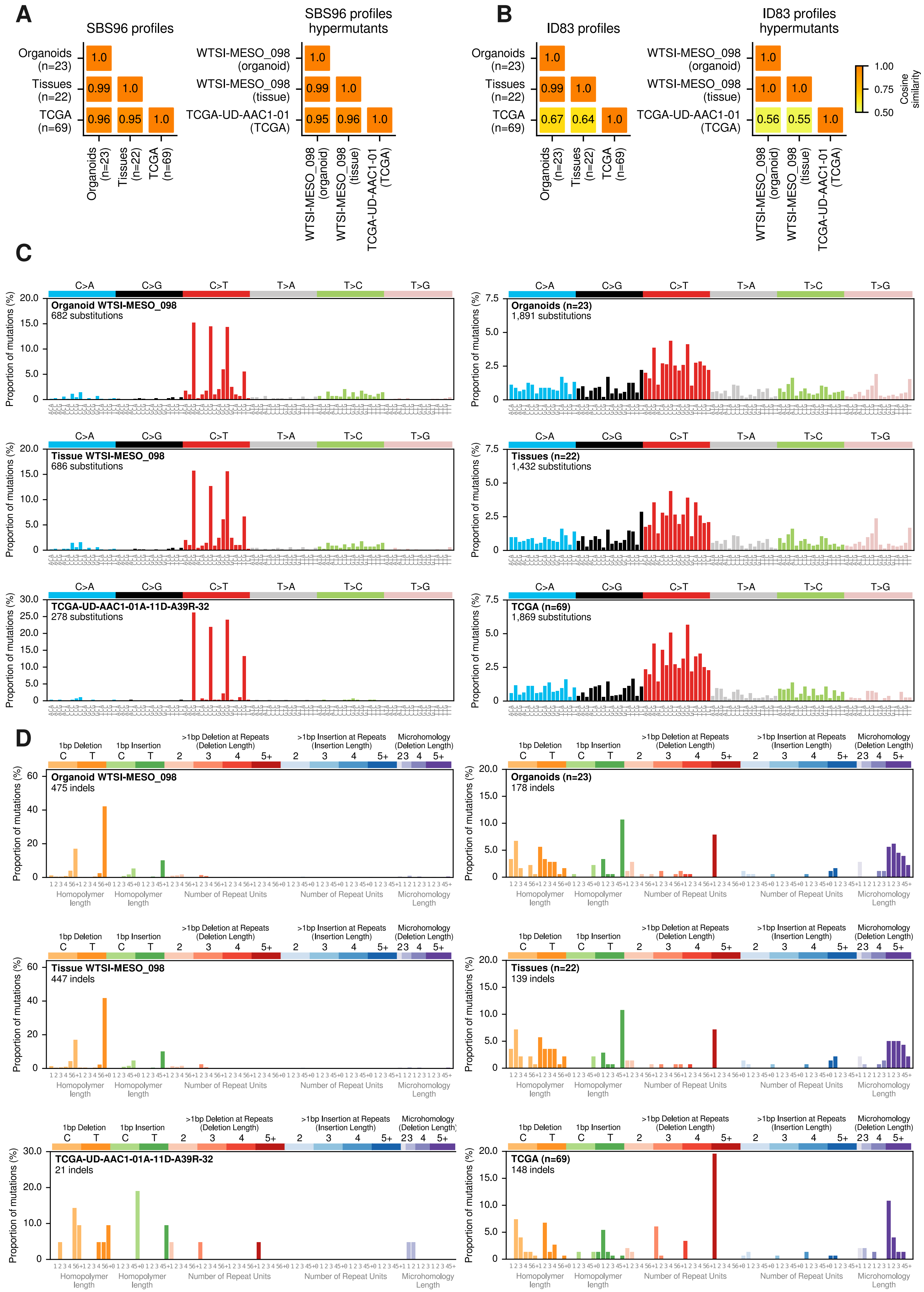

#### Extended Figure E4

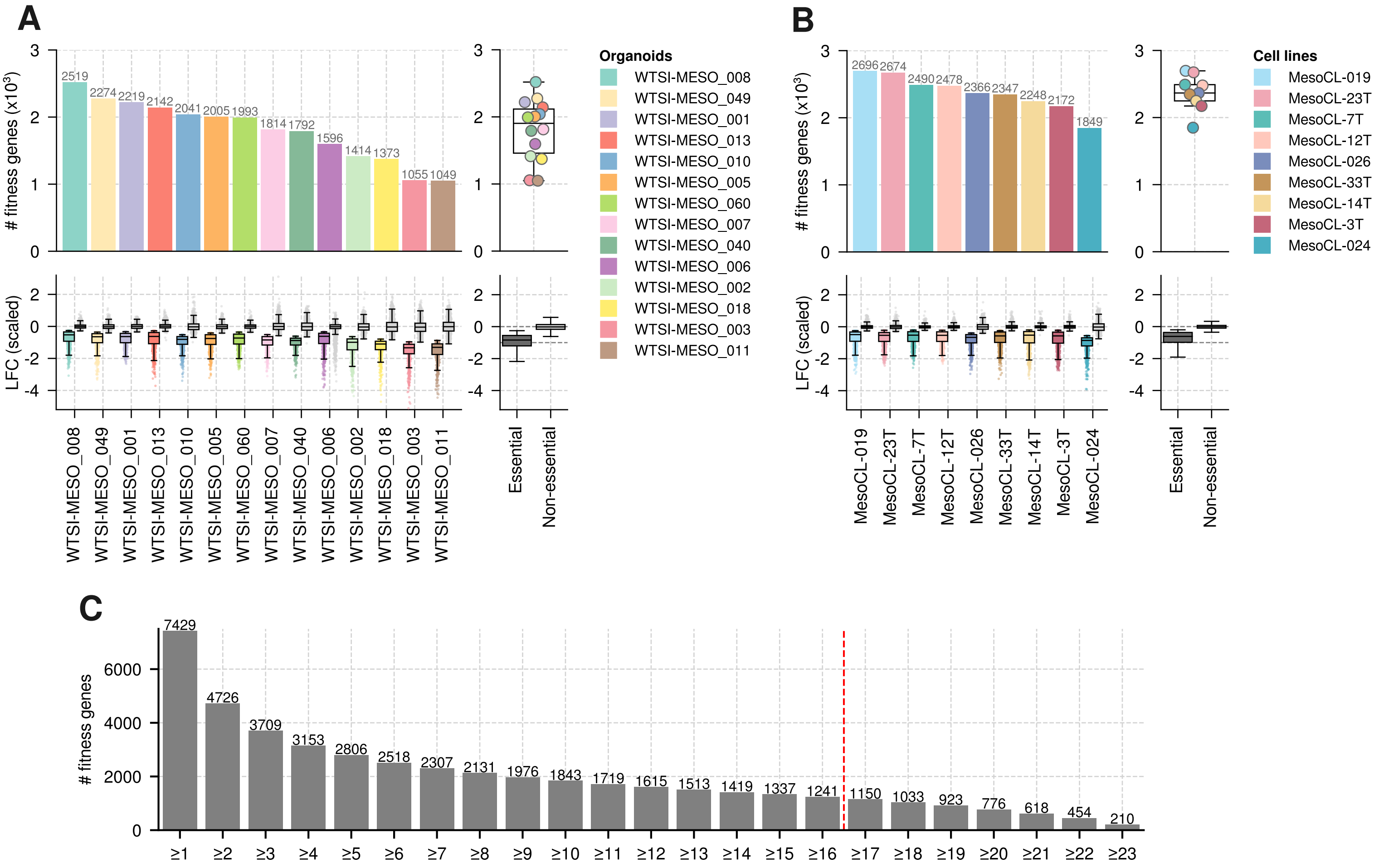
